# Myc instructs and maintains pancreatic adenocarcinoma phenotype

**DOI:** 10.1101/556399

**Authors:** Nicole M. Sodir, Roderik M. Kortlever, Valentin J.A. Barthet, Luca Pellegrinet, Tania Campos, Steven Kupczak, Lamorna Brown Swigart, Laura Soucek, Mark J. Arends, Trevor D. Littlewood, Gerard I. Evan

## Abstract

Pancreatic ductal adenocarcinoma (PDAC) is characterized by its dismal prognosis and its signature fibroinflammatory phenotype. We show that activation of Myc in PanIN epithelial cells is alone sufficient to instruct and maintain immediate transition of indolent PanINs to PDACs phenotypically identical to the spontaneous human disease. Myc does this by inducing a distinct, tissue-specific ensemble of instructive signals that, together, coordinate changes in multiple, stromal and inflammatory cell types to generate the signature PDAC stroma. We also demonstrate that the Myc PDAC switch is completely reversible and that Myc deactivation immediately triggers meticulous disassembly of both PDAC tumor and stroma. Hence, both the formation and deconstruction of the complex PDAC phenotype may be mediated by a single, reversible molecular switch.

**SIGNIFICANCE:** Pancreatic ductal adenocarcinoma (PDAC) has a dismal prognosis and lacks effective therapies. We show that Myc is a single molecular switch that directly and immediately instructs transition from indolent KRas^*G12D*^-induced PanIN to the characteristic complex, multi-cell-type fibroinflammatory and immune-cold PDAC phenotype through the release of a distinct, tissuespecific set of instructive signals. The same combination of KRas^*G12D*^ and Myc drives a very different phenotype in lung, indicating that the principal phenotypes of adenocarcinomas are dictated by tissue of origin not specific oncogenes. We also show that the Myc switch is immediately and completely reversible: blocking Myc function triggers meticulous disassembly of the entire PDAC tumor-stromal edifice demonstrating that phenotypic complexity is not a barrier to effective treatment of cancers.

## INTRODUCTION

Pancreatic ductal adenocarcinoma (PDAC) is a lethal cancer with a 5-year survival of less than 5% ^1^. Its dismal prognosis is in great part due to its typically late presentation and its profound resistance to all forms of conventional chemo and radiotherapy. Studies of resected tumors suggest that malignant ductal adenocarcinoma evolves from pancreatic intraepithelial neoplasms (PanIN), indolent precursors that are histologically classified into four stages (PanIN1A, 1B, 2 and 3) based on increasing degrees of nuclear and architectural atypia ^2-4^. Frank PDAC is characterized by its striking fibroinflammatory stroma, absent from normal pancreas, whose attendant dense desmoplasia typically constitutes some 90% of tumor bulk ^8^,^9^. This fibroinflammatory stroma is a product of complex interplay between tumor cells and adjacent mesenchymal, endothelial, inflammatory and immune cells and is thought to contribute to PDAC therapeutic recalcitrance by impeding vascular perfusion and oxygenation ^10,11^.

At the molecular level, PDAC is associated with a core of recurring, presumably causal, oncogenic mutations, of which activating mutations in KRas appear to be founder drivers and are present in over 90% of both PanINs and PDAC. Other frequently recurring signature mutations inactivate *CDKN2A* (90–95%), *TP53* (50–84%) or *SMAD4/DPC4* (49–55%) and recent, more granular, analyses reveal a plethora of additional sporadic PDAC mutations encompassing great functional diversity ^5-7^. As the most commonly recurring oncogenic mutation, many of the canonical features of PDAC are attributed to precocious KRas signaling. KRas is a multifunctional communication node with diverse, pleiotropic outputs that encompass downstream intracellular effectors (e.g. MAPK, PI3K, p38, JAK/STAT, Hippo and NF-κB) and a host of extracellular signals and their receptors (e.g. Wnt, Notch, Shh, CXCL1, 2 and 5, CXCL8 (IL-8), IL-6, GM-CSF and IL-17RA). These influence diverse aspects of PDAC pathology, including tumor initiation, maintenance, drug sensitivity, metabolism, macropinocytosis, metastasis, desmoplasia, inflammation and immunity ^12,13^. Oncogenic KRas has been successfully used as the initiating driver in various pancreas cancer mouse models ^14-22^. All concur that, without cooperating oncogenic mutations, KRas oncogenic potential is very poor, stalling at indolent PanIN with little desmoplasia, inflammation or lymphocytic suppression ^14,23^. Aside from its role in pancreatic tumor initiation (Collins, 2014 #433), oncogenic KRas also plays an obligate role in maintenance of established PDAC ^14,16,23,24^ and hence appears to provide a necessary platform upon which subsequent cooperating lesions exert their oncogenic impacts.

The archetypical cooperating oncogenic partner of KRas is Myc ^25^, a pleiotropic transcription factor whose putative role is to coordinate expression of thousands of genes whose products have enormously diverse functions involved in virtually every aspect of cell and tissue behavior, that together mediate somatic cell proliferation. In proliferating normal cells, Myc serves as a functionally non-redundant and essential transcriptional node that is tightly controlled by mitogen availability ^26^. By contrast, some degree of aberrant Myc regulation is implicated in the majority, maybe all, cancers. Such aberrant Myc expression only seldom involves mutation of the *myc* gene itself and is usually an indirect consequence of its relentless induction by “upstream” oncogenes – notable examples being Wnt, Notch and Ras itself ^27-33^. During pancreas organogenesis and early postnatal growth, Myc is highly expressed in a population of multipotent pancreatic progenitor cells and is, like its immediate upstream activators β-catenin ^33-35^ and Notch ^32^, required for exocrine pancreas development, maturation and neonatal expansion ^31,36-39^. While Myc is not appreciably expressed in normal adult, uninjured, non-proliferating pancreas, >40% of PDACs exhibit marked expression of Myc and its canonical target gene signature while overt amplification of the *MYC* gene at 8q24 is frequent in especially aggressive tumors ^19,31,40-42^. Furthermore, multiple *in vivo* mouse studies indicate that deregulated Myc is causally oncogenic in exocrine pancreas (reviewed in ^31^). Thus, classical transgenic over-expression of Myc targeted to mature acinar cells via an elastase promoter induces widespread acinar-to-ductal metaplasia (ADM) ^10^ while *pdx-1* promoter-directed over-expression of Myc in pancreatic progenitor cells through organ development elicits rapid emergence of ductal PanIN ^43^. Nonetheless, such tumors only rarely progress to PDAC ^44^, in part because high levels of Myc trigger potent tumor suppressor responses ^45^. Conversely, *pdx-1*-CRE-driven excision of endogenous *myc* concurrently with KRas^*G12D*^ activation blocks transition of PanINs to PDAC. Studies also indicate a persistent maintenance role for Myc in established PDAC. *myc* knockdown inhibits proliferation of human PDAC cell lines *in vitro* ^46^ and direct ^43^ or indirect ^47^ inactivation of Myc triggers macroscopic regression of established PDAC in mouse models *in vivo* ^48^, albeit with persistence of dormant tumor cells ^43^. Taken together, these data suggest a critical role for Myc in both the progression from PanIN to PDAC and in subsequent PDAC maintenance ^32^. However, the nature of this role remains obscure. For example, Myc has no obvious functional overlap with any of the canonical mutations in *CDKN2A, TP53* and *SMAD4* that typically cooperate with KRas in PDAC.

Here we use a rapidly and reversibly switchable genetic mouse model whereby Myc may be engaged or disengaged, at will, in indolent KRas^*G12D*^-induced PanINs to address key questions concerning the mechanistic role of Myc in PDAC. Does Myc activation drive immediate transition of PanIN to PDAC or, instead, merely raise the eventual likelihood that sporadic progression eventually occurs? What is the phenotype of the pancreas tumors that Myc activation elicits and do they resemble spontaneous PDACs? Finally, we address why Myc is required for the maintenance of PDAC tumors and by what mechanism PDAC tumors regress upon Myc deactivation.

## RESULTS

### Myc deregulation drives immediate progression of KRas^G12D^-induced PanINs to adenocarcinoma

Conditional expression of oncogenic KRas^*G12D*^ in pancreas progenitor cells drives protracted and sporadic outgrowth of indolent PanIN lesions ^17^. Rarely, these progress to overt adenocarcinoma through aleatory accretion of additional lesions. Since deregulated and/or over-expressed Myc is a frequent feature of advanced PDAC, we first asked whether Myc deregulation is sufficient to drive progression of indolent KRas^*G12D*^-driven PanINs to PDAC using the well-characterized *pdx1-Cre;LSL-KRas*^*G12D/+*^ model (*K*^*pdx*1^) in which Cre recombinase expression, driven by the *pdx/IPF1* (*Pancreas/duodenum homeobox protein 1*) promoter in embryonic pancreatic and duodenal progenitor cells from around E8 ^18^, triggers expression of a single allele of KRas^*G12D*^ driven from the endogenous *KRas2* promoter. *K*^*pdx*1^ mice with a homozygous *Rosa26-LSL-MycER*^*T*2^ ^45^ background were generated. In the resulting *pdx1-Cre;LSL-KRas*^*G12D*^*;Rosa26LSL-MycER*^*T*2^ (*KM*^*pdx1*^) mice, activation of Cre recombinase in pancreatic progenitor cells induces expression of both KRas^*G12D*^, driven from its endogenous promoter, and the conditionally regulatable MycER^T2^ variant of c-Myc, driven constitutively from the promiscuously active *Rosa26* promoter at quasi-physiological levels. Although expressed constitutively in *KM*^*pdx*1^ pancreatic epithelia, MycER^T2^ is active only in the presence of its ligand, 4-hydroxytamoxifen (4-OHT).

As previously described ^17^, by 15 weeks of age *K*^*pdx*1^ control mice (i.e. KRas^*G12D*^ alone) each exhibited a small number of early-stage PanIN-1A and PanIN-1B lesions, with the vast majority of pancreatic ducts appearing histologically normal. These PanIN lesions were small, with little or no desmoplastic stroma (Figure 1A). Activation of MycER^T2^ alone for 3 weeks in the absence of KRas^*G12D*^ (*M*^*pdx*1^ mice) had no significant histological impact (Figure 1A) save for a very modest increase in proliferative index (data not shown). By contrast, sustained activation of MycER^T2^ for 3 weeks in *KM*^*pdx*1^ mice (i.e. KRas^*G12D*^ and Myc together) induced dramatic progression of existing PanINs to invasive adenocarcinomas that bore all the stereotypical features of spontaneous mouse and human PDAC: highly proliferative nests of epithelial tumor cells embedded in extensive avascular, hypoxic, αSMA (α-smooth muscle actin)-positive desmoplastic stroma densely infiltrated with leukocytes, yet largely devoid of T lymphocytes (Figures 1B, 7, S1 and S8). Tamoxifen alone had no discernible impact on pancreas architecture (Figures 1B and S1). Identical observations were made using the complementary *p48-Cre;LSL-KRas*^*G12D*^*; Rosa26LSL-MycER*^*T*2^ (*KM*^*p*48^) mouse PDAC model, in which Cre expression is driven slightly later and in more pancreas-specific progenitors by the *p48/PTF1* promoter (Figure S1) ^17,49^.

**Figure 1.**
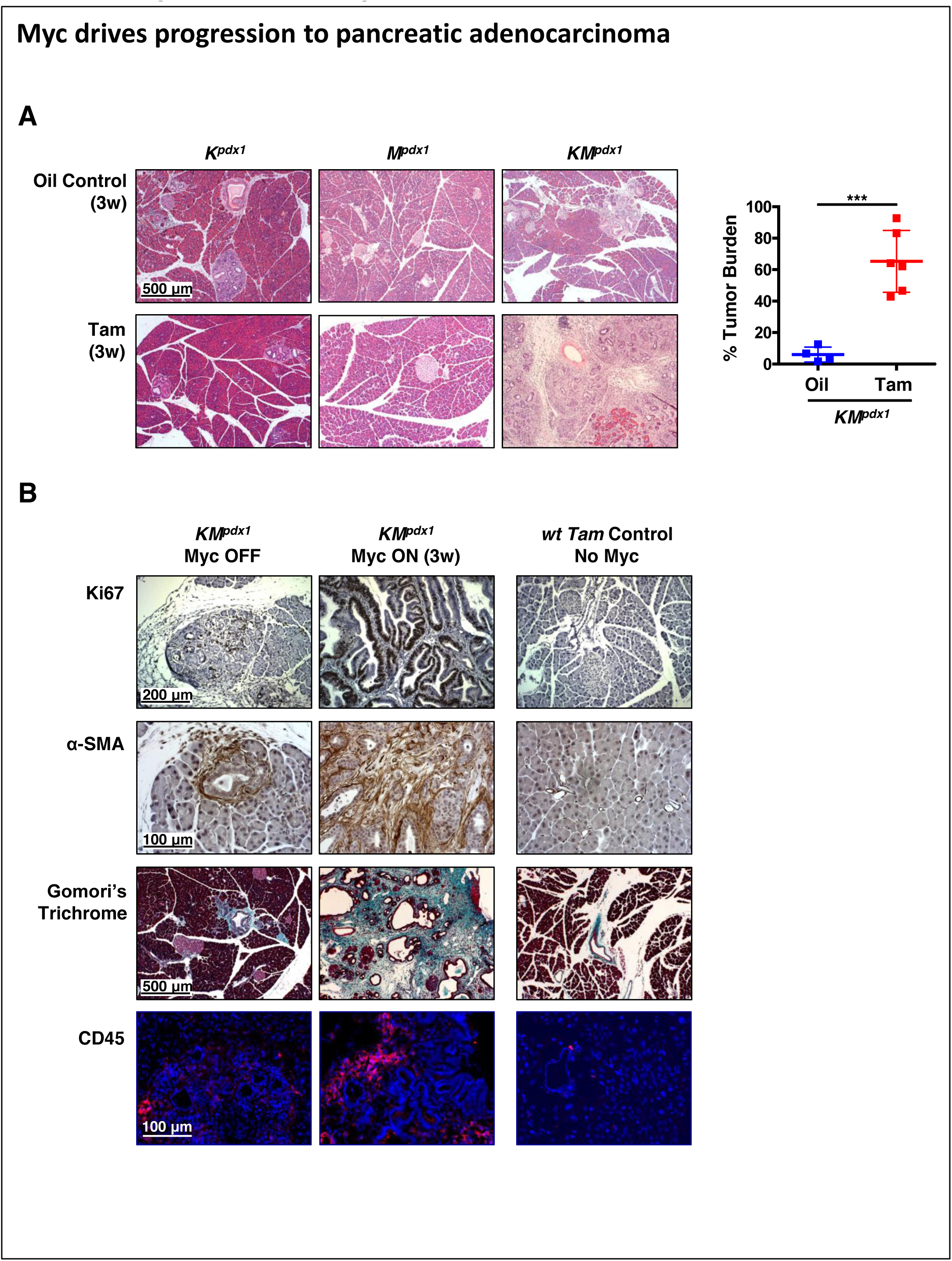
Myc drives progression to pancreatic adenocarcinoma (related to Figures S1, S3 and S4) (A) Representative H&E stained sections of pancreata from 15-week old *K*^*pdx*1^ (Kras^G12D^ only; left), *M*^*pdx*1^ (MycER^T2^ only; middle), and *KM*^*pdx*1^ (MycER^T2^ and Kras^G^12^D^; right) mice treated with either oil control (top row) or tamoxifen (Tam) (bottom row) for 3 weeks. Percentage tumor burden relative to the total pancreas from *KM*^*pdx*1^ mice treated with either oil (control) or tamoxifen is shown right. Results represent mean ± SD, n = 4-6 mice per treatment group. The unpaired Student’s t-test was used to analyze tumor burden. ***p < 0.001. SD = standard deviation. (B) Representative immunohistochemical staining (IHC) for proliferation (Ki67, top row), α-smooth muscle action (α-SMA) (second row), collagen deposition by Gomori’s Trichrome staining (blue-stained collagen, third row) and immunofluorescence staining (IF) of CD45 expressing leukocytes (red, bottom row) in sections of pancreata harvested from 15 week old *KM*^*pdx*1^ mice treated with either oil (Myc OFF, left) or tamoxifen (Myc ON, middle) for 3 weeks and wild type control mice treated with tamoxifen (No Myc, right).

Although expression of MycER^T2^ is (like KRas^*G12D*^) restricted solely to epithelial cells of *KM*^*pdx*1^ and *KM*^*p*48^ tumors, the stereotypical fibroinflammatory and hypovascular stromal changes that Myc induces involve diverse stromal hematopoietic, endothelial and mesenchymal cell types. To address whether the Myc-induced PDAC phenotype is a direct, instructive result of Myc action or, instead, a passive chronic consequence of constraints by the host pancreatic tissue we made use of the inherent rapidity and synchrony of MycER^T2^ activation by tamoxifen *in vivo* ^50-53^. Myc was activated in *KM*^*p*48^ mice by systemic administration of tamoxifen for 0, 1 or 3 days, and pancreata harvested and analyzed. Within 24 hours of Myc activation, we observed a dramatic increase in proliferation of the MycER^T2^-expressing epithelial tumor compartment of all PanIN lesions and this was accompanied by diverse stromal changes: most notably a marked influx of F4/80^+^ macrophages, Ly-6B.2^+^ neutrophils and B220^+^ B lymphocytes, concurrent loss of CD3^+^ T cells and induction of αSMA in proximal stellate cells, indicative of myofibroblastic transdifferentiation (Figure 2). Rather later (clearly evident by 72 hours) we observed progressive deposition of dense desmoplasia, loss of patent blood vessels and induction of HIF1-α, a marker of hypoxia (Figure 2), all expected secondary consequences of prior stellate cell activation. Identical Myc-induced phenotypic changes, with identical kinetics, were observed in pancreata of the complementary *KM*^*pdx*1^ switchable Myc model (Figure S2). Hence, all the signature tumor-stromal features of aggressive spontaneous PDAC seen in spontaneous mouse and human PDAC are instructed by activated Myc from the outset.

**Figure 2.**
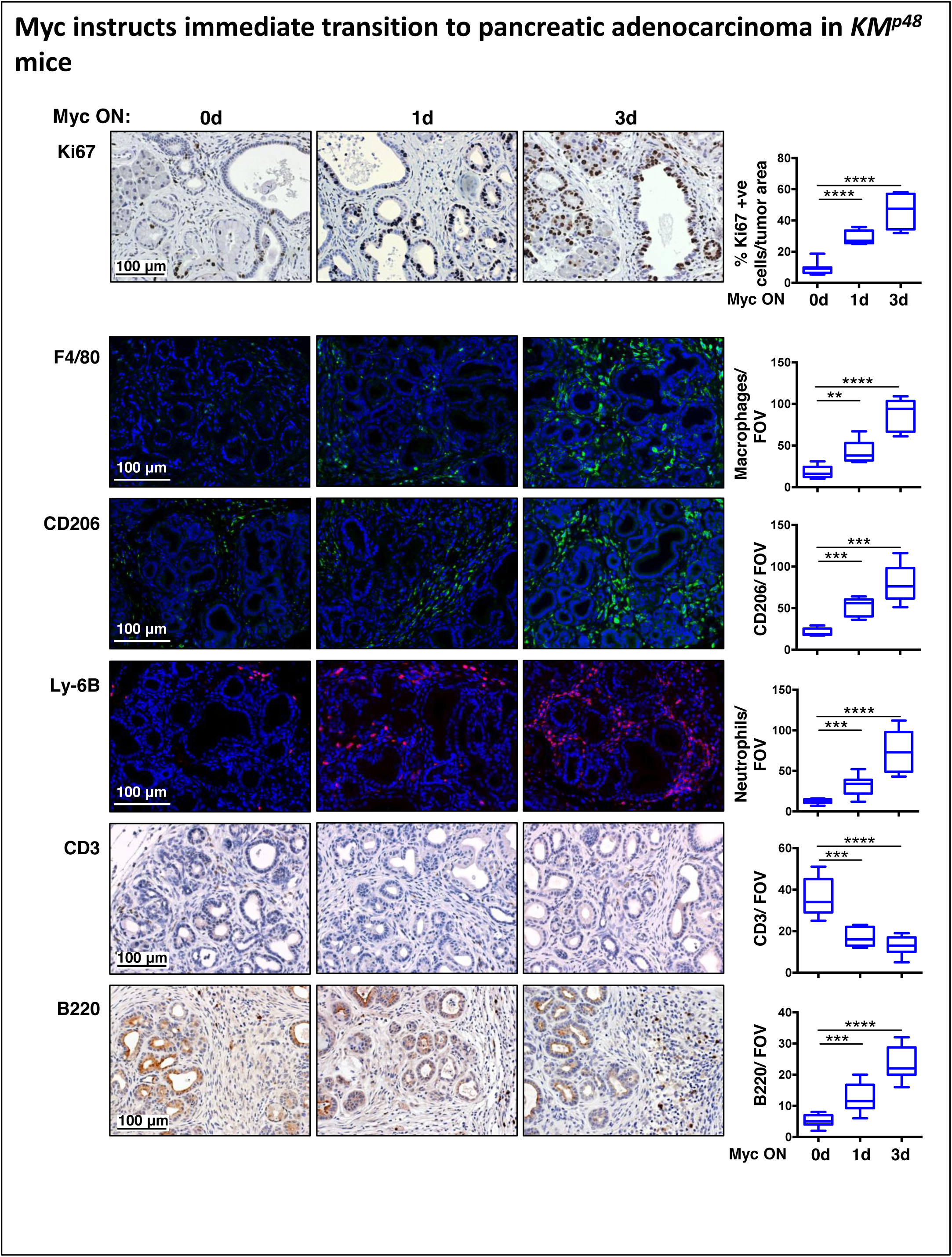
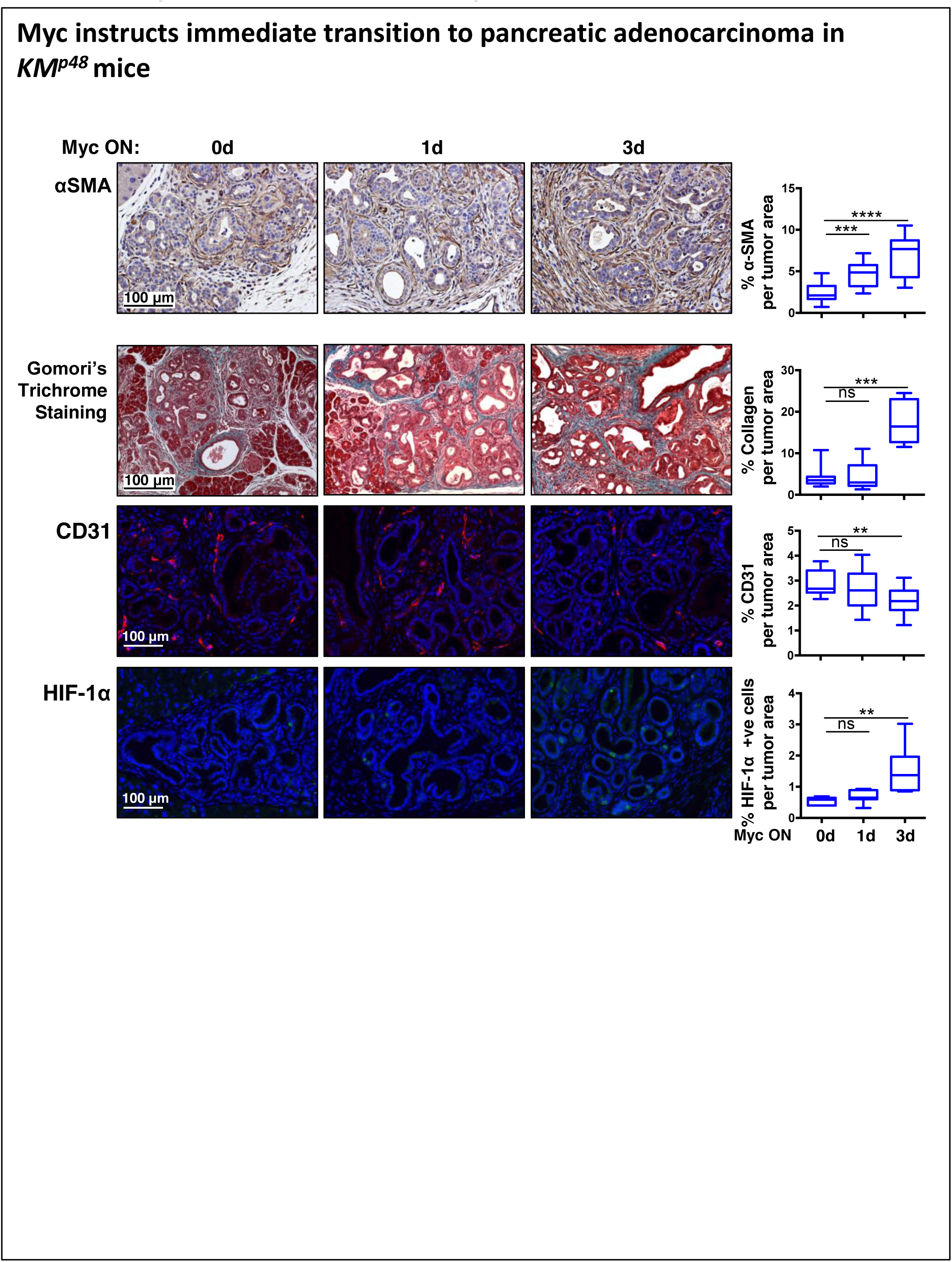
Myc instructs immediate transition to pancreatic adenocarcinoma in *KM*^*p48*^ mice (Related to Figure S2) Immunohistochemical staining (IHC) for proliferation (Ki67), immunofluorescence (IF) staining of macrophages (F4/80, CD206) and neutrophils (Ly-6B), IHC for T cells (CD3), B220^+^ lymphocytes and α-SMA, Gomori’s Trichrome staining of collagen, and IF for vasculature (CD31) and hypoxia (HIF-1α) in sections of pancreata harvested from 12 week old *KM*^*p*48^ mice treated with oil (left panel) or tamoxifen for 1 or 3 days (Myc ON). Scale bars apply to each row. Quantification (Box-and whisker plots, right) shows upper extreme, upper quartile, median, lower quartile, lower extreme. n = 4-6 for each treatment group. **p < 0.01, ***p < 0.001, ****p < 0.0001 (ns = non-significant). Data were analyzed using unpaired t-test. FOV = field of view.

Unexpectedly, sustained activation of MycER^T2^ proved rapidly lethal in *KM*^*pdx*1^ mice but not in their *KM*^*p*48^ counterparts. Such lethality corresponded with extensive Myc-induced overgrowth of *KM*^*pdx*1^ duodenal epithelium, severely disrupted crypt-villus architecture and multiple intestinal neoplasms (Figure S3). This is likely due to the known activity of the *pdx-1* promoter in primordial duodenum - a promiscuity not shared by the *p48* promoter, whose activity is tightly restricted to pancreatic precursor cells ^17^. However, since the PDAC phenotype induced by Myc was identical in *KM*^*pdx*1^ and *KM*^*p*48^ pancreata, all long-term Myc activation studies were thenceforth conducted in *KM*^*p*48^ mice. Extended activation of MycER^T2^ eventually also proved lethal in *KM*^*p*48^ animals, however, due to the outgrowth of grossly expanded, highly desmoplastic PDAC (Figure S4): *KM*^*p*48^ mice retained normal intestinal morphology throughout, albeit punctuated by frequent PDAC metastases (data not shown).

### Specific signals instruct the Myc-induced PDAC phenotype

MycER^T2^ is expressed only in the epithelial compartments of both *KM*^*pdx*1^ and *KM*^*p*48^ pancreas tumors. Hence, the immediate stromal changes that Myc elicits must be instructed by signals originating in those MycER^T2^-expressing tumor epithelial cells. To identify such signals, we activated MycER^T2^ in *KM*^*p*48^ mice with a single systemic dose of tamoxifen, harvested pancreata 6 hours later, and then interrogated tissue extracts with growth factor/cytokine arrays and isolated mRNA by RT-PCR. Several discrete growth factors and regulators of inflammatory and immune cells were induced (Figure 3), including SHH, previously shown to be critical for PDAC desmoplastic deposition and maintenance ^11^.

**Figure 3:**
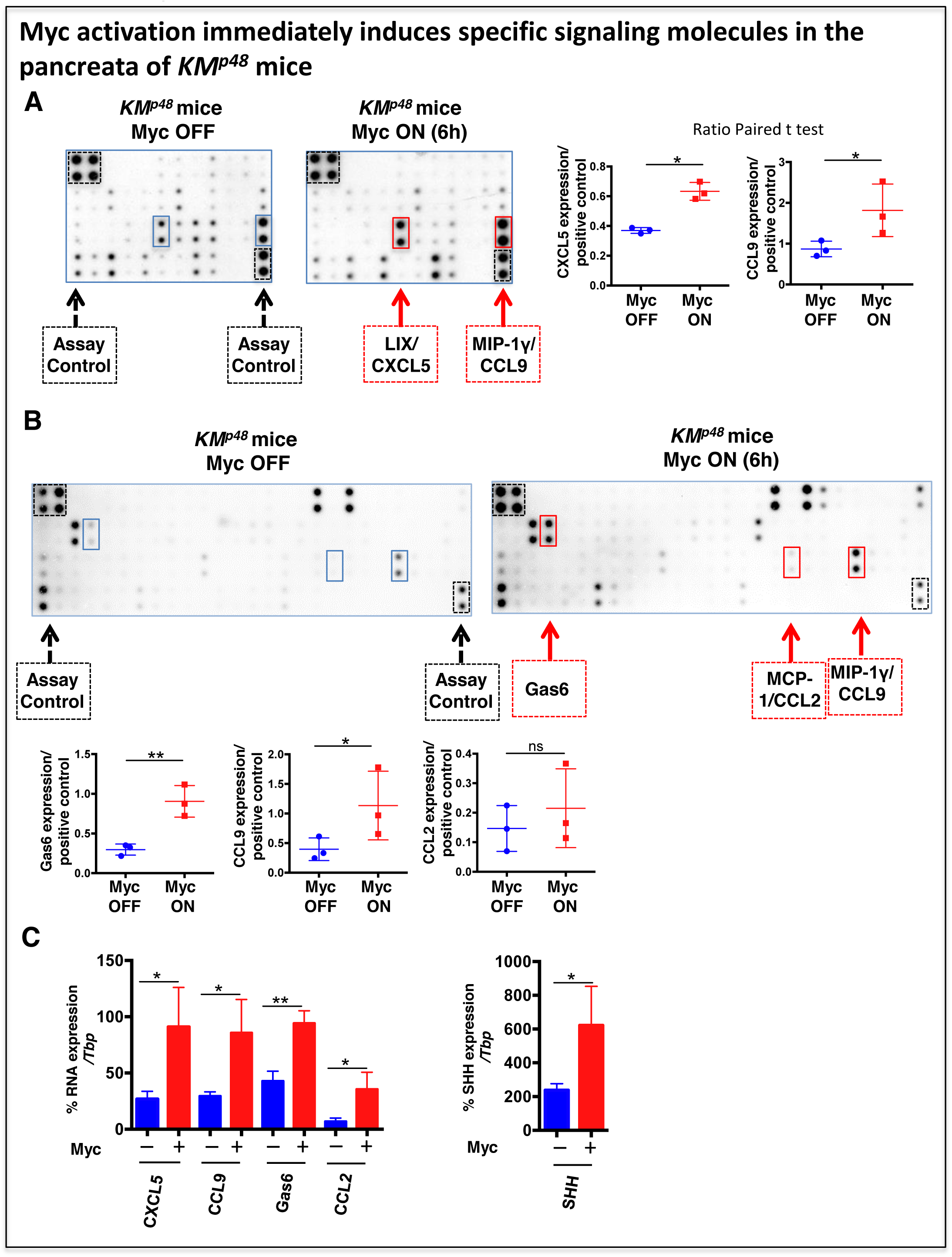
Myc activation immediately induces specific signaling molecules in the pancreata of *KM*^*p*48^ mice. (A) and (B) Representative inflammation antibody arrays of 40 targets (A) and cytokine antibody arrays of 97 targets (B) probed with whole pancreas protein lysates from 12-week old *KM*^*p*48^mice treated for 6 hours with oil (Myc OFF, left) or tamoxifen (Myc ON, right). Fold change quantitation of CXCL5, CCL9, Gas6 and CCL2 signals derived from protein arrays are shown. Each data point represents a single mouse. n = 3 for each cohort. Ratio paired t-test is used to analyze the data. *p < 0.05, **p < 0.01, ns = not significant. (C) Quantitative RT-PCR for *CXCL5, CCL9, GAS6, CCL2* and *SHH* mRNA derived from 12-week old *KM*^*p*48^ pancreata treated for 6 hours with oil (blue bars) or tamoxifen (red bars). *Tbp* was used as an internal amplification control. The unpaired t-test with was used to analyze Taqman expression data. n = 3 for each cohort. The mean ± SD is shown. *p < 0.05, **p < 0.01.

Macrophage influx into PanINs, clearly evident by 24 hr post Myc activation, was presaged by induction at 6 hr of two well-documented macrophage chemoattractants, CCL9 (MIP1γ) and CCL2 (MCP1), the latter widely implicated in human PDAC ^55^. Myc-induced influx of Ly-6B+ neutrophils was preceded at 6hr by induction of CXCL5 (Figure 3), a well-characterized neutrophil attractant ^59,60^ and one of several CXCR2 (IL-8RB) receptor ligands. We also observed Myc-dependent upregulation of GAS6 (Figure 3), a ligand for AXL kinase that has previously been implicated in diverse neutrophil dynamics ^61-63^. Moreover, GAS6 induction correlated with appearance of pAXL immunoreactivity in a portion of PanIN epithelial cells (Figure 4A) while blockade of either GAS6, using low dose Warfarin ^64,65^ (Figure 4B), or of AXL kinase, using a specific inhibitor TP-0903 ^66^ (Figure 4C), profoundly inhibited Myc-induced neutrophil influx into PanINs, significantly retarded Myc-driven stellate cell activation and proliferation (Figure S5) and impeded tumor growth. Surprisingly, AXL inhibition did not discernibly inhibit Myc induction of CXCL5 (data not shown). Hence, GAS6 and AXL are components of a novel, obligate Myc-driven neutrophil recruitment pathway in PDAC. Unexpectedly, AXL inhibition also profoundly blocked Myc-induced peritumoral influx of B220+ B lymphoid cells but only modestly inhibited Myc-induced macrophage influx (Figure 4C). AXL inhibition also had no discernible impact on the rapid depletion of CD3^+^ T cell that Myc induces (data not shown). However, we recently showed that Myc-induced depletion of CD3+ T cells from lung adenomas is dependent upon up-regulation of PD-L1 (CD274/B7-H1), a ligand for the PD1 (CD279) receptor whose ligation quells T lymphocyte immune functions. Immunohistological analysis revealed that Myc activation rapidly induced PD-L1 expression on both PanIN epithelial cells *in vivo* (Figures 5A and S6A) and on isolated *KM*^*pdx*1^ PDAC cells *in vitro* (Figure S6B). Moreover, ChIP analysis of cultured *KM*^*pdx*1^ PDAC cells located MycER^T2^ protein on the *CD274* (*PD-L1*) gene (Figure S6C), indicating that Myc-dependent PD-L1 induction in PanIN epithelium is cell autonomous and the *B7-H1* gene is most probably a direct Myc target. As in lung, systemic blockade of PD-L1 completely abrogated Myc-induced T cell depletion in pancreas tumors (Figure 5B and 5C). However, as with lung, PD-L1 blockade and consequent persistence of CD3^+^ T cells had no discernible inhibitory impact on subsequent tumor growth (data not shown). In summary, we identify multiple candidate signaling molecules responsible for the diverse, tightly coordinated transition from indolent PanIN to aggressive PDAC that Myc induces.

**Figure 4.**
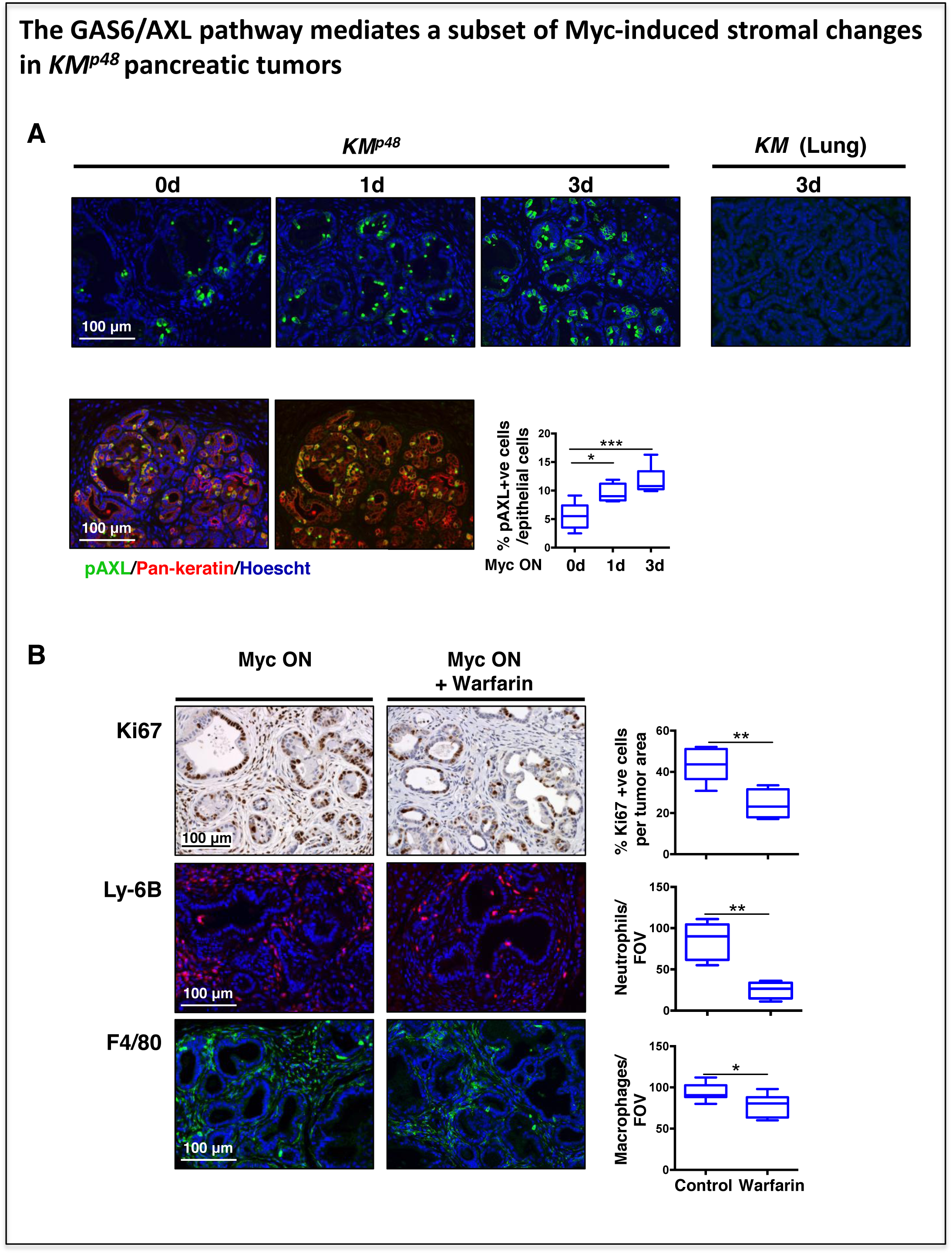
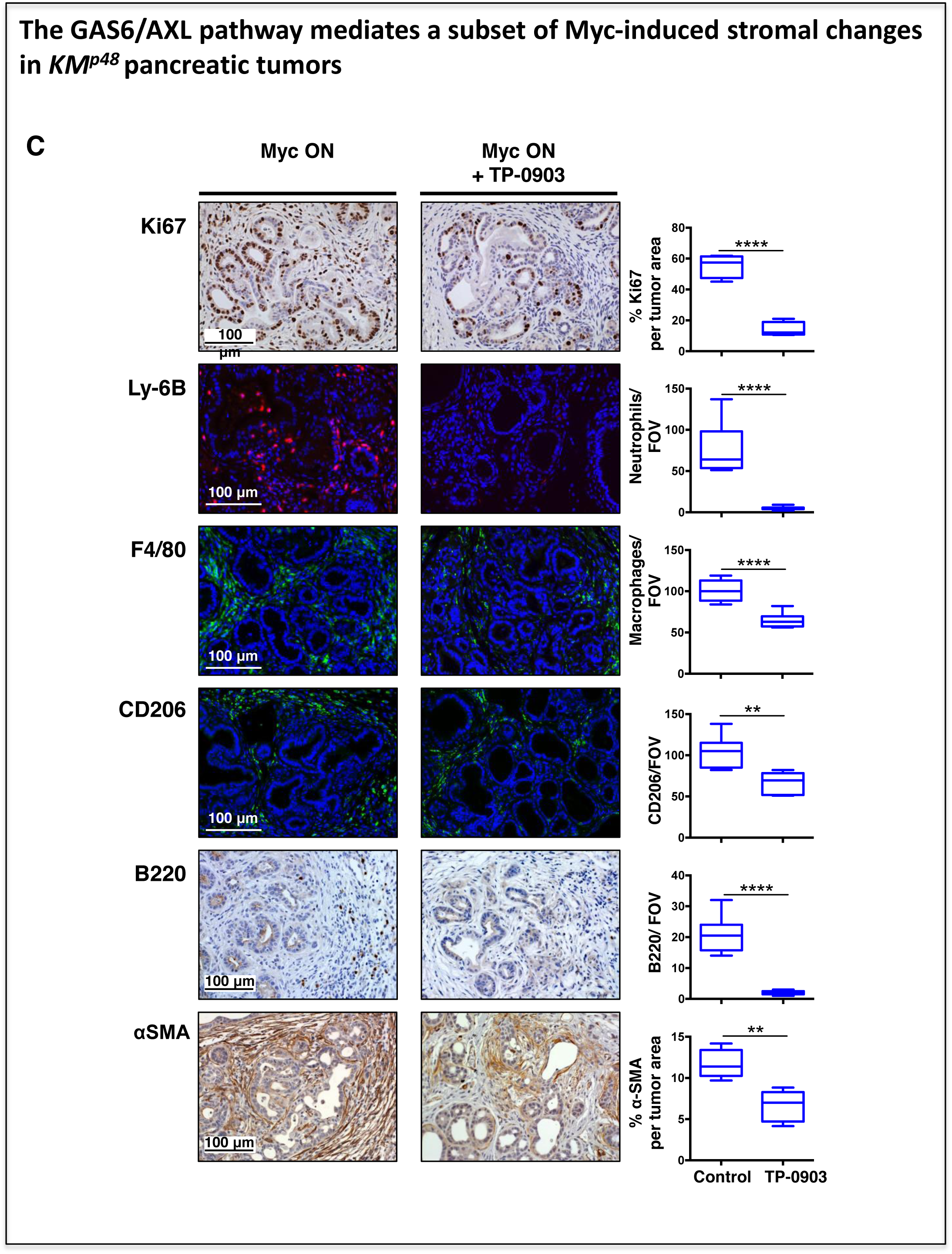
The GAS6/AXL pathway mediates a subset of Myc-induced stromal changes in *KM*^*p*48^ pancreatic tumors (related to Figure S5) (A) Immunofluorescence staining of phospho-AXL (pAXL) (green) and its quantification in sections of pancreata of 12-week old *KM*^*p*48^ mice treated with oil (Myc OFF, top left) or tamoxifen for 1 (middle) or 3 days (right) (Myc ON) and of lungs from Adeno-Cre treated *kRas*^*G12D/+*^*; MycER*^*T2*/*T2*^ mice treated with tamoxifen (Myc ON) for 3 days (extreme right). Representative co-localization of pAXL (green) and pan-keratin (red, epithelial cells) (bottom row). Scale bars apply across each row. (B) Representative immunohistochemical staining for proliferation (Ki67, top panel) and immunofluorescence staining of neutrophils (Ly-6B, middle) and macrophages (F4/80, bottom) in sections of pancreata harvested from 12-week old *KM*^*p*48^ mice either without (left) or with warfarin for 4 days commencing one day prior to tamoxifen administration (Myc ON). Quantitation is shown on the right. n = 4 - 6 per group. *p<0.05, **p < 0.01 (C) Representative immunohistochemical or immunofluorescence staining of proliferation (Ki67), neutrophils (Ly-6B), macrophages (F4/80, CD206), B220 lymphocytes and α-SMA of pancreata harvested from 12-week old *KM*^*p*48^ mice untreated (left) or treated (right) with pAXL inhibitor TP-0903 concurrent with tamoxifen administration (Myc ON) for 5 days. Scale bars apply across each row. Quantitation is shown on the right. n = 4 - 6 per group. *p<0.05, **p < 0.01, ***p < 0.001, ****p < 0.0001. FOV = field of view.

**Figure 5.**
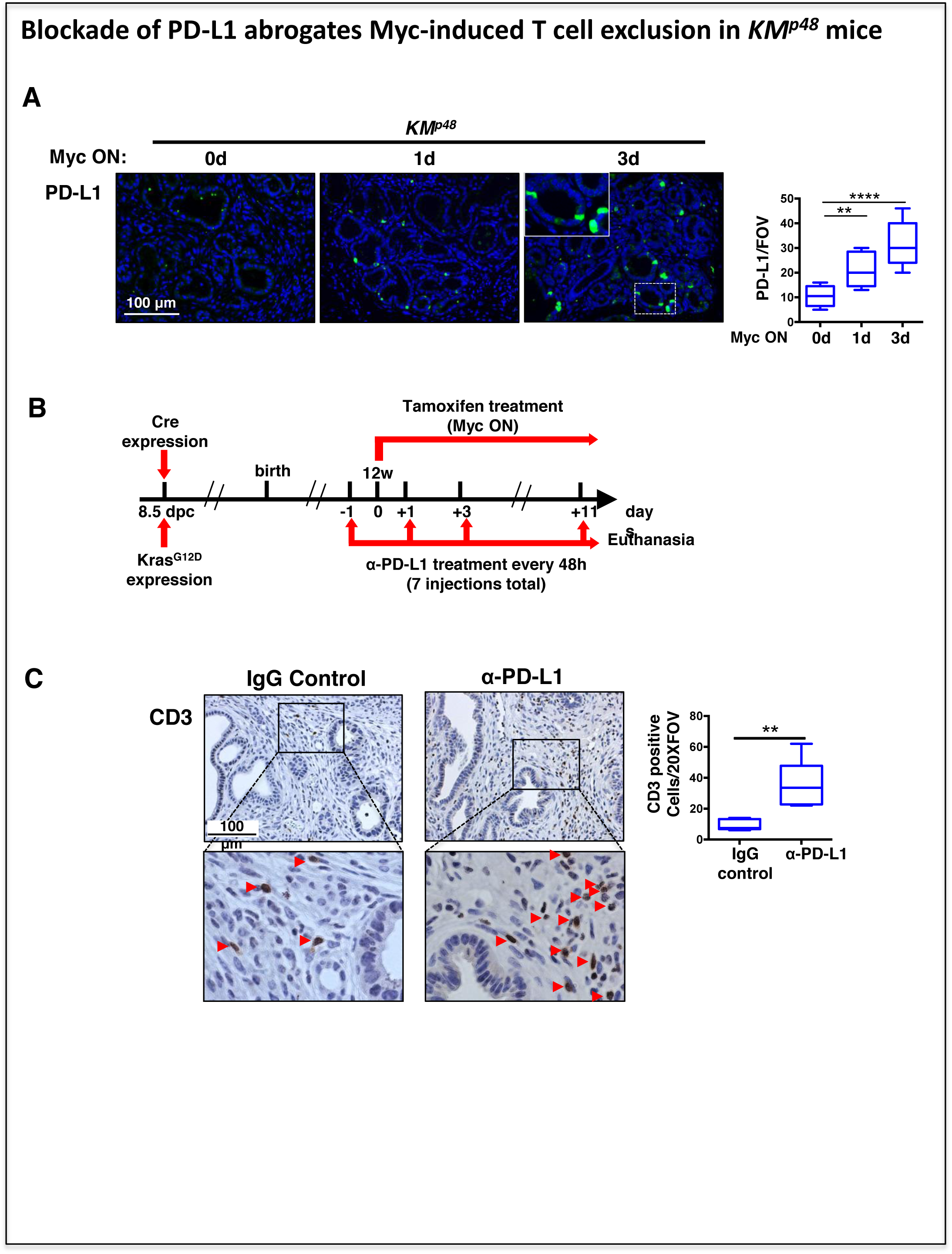
Blockade of PD-L1 abrogates Myc-induced T cell exclusion in *KM*^*p*48^ mice (related to Figure S6) (A) Immunofluorescence staining of PD-L1 in sections of pancreata harvested from 12-week old *KM*^*p*48^ mice treated with oil (Myc OFF, left) or tamoxifen (Myc ON) for 1 (middle) or 3 days (right). Upper boxed region shown at higher magnification. Proportion of cells expressing PD-L1 is shown on the right. n = 4–6 per treatment group. FOV = field of view. (B) Schematic representation of PD-L1 blocking experiment: starting 24 hours prior to tamoxifen administration (Myc ON), 12-week old *KM*^*p*48^ mice were injected intraperitoneally every two days for two weeks with neutralizing antibodies for PD-L1 or IgG control. (C) Representative immunohistochemical staining of T cells (CD3) in pancreatic sections from mice described in (B). Boxed regions are shown at higher magnification below. Red arrows indicate CD3-positive cells. Quantitation of CD3-positive cells shown right. n = 4–6 per treatment group. FOV = field of view. The unpaired student t-test was used to analyze data. n = 4 - 6 per group. **p < 0.01, ****p < 0.0001.

### The Myc-driven transition from PanIN to adenocarcinoma is immediately and fully reversible

We next addressed whether persistent Myc activity is required to maintain PDAC. MycER^T2^ was activated for 3 weeks in 12-week-old *KM*^*pdx*1^ or *KM*^*p*48^ animals, inducing widespread multifocal aggressive PDAC in all mice. Subsequent MycER^T2^ de-activation triggered complete regression of all adenocarcinomas (Figure 6A and data not shown), culminating in pancreata largely indistinguishable from those in which Myc had never been activated. Even grossly sized pancreatic tumors, generated after 4-6 weeks MycER^T2^ sustained activation, regressed with identical histology and kinetics following Myc de-activation (Figures S7A and S7B), although these often exhibited abnormalities in gross organ architecture with evidence of extensive scarring and numerous cyst-like structures (Figure S7C). To confirm that such Myc dependency is not a property peculiar to pancreatic adenocarcinomas induced by direct Myc activation, we also blocked endogenous Myc activity in spontaneously evolving KRas^*G12D*^-driven PDAC using the Omomyc dominant negative Myc dimerization mutant (Figure 6B). To do this, we used *pdx1-Cre;LSL-KRas*^*G12D*^*; p53ER*^*TAM*^ *KI;TRE-Omomyc;CMVrtTA* (*KPO*) mice in which *pdx1-Cre-*driven activation of oncogenic KRas^*G12D*^ is combined with a background in which the endogenous p53 of *KPO* mice is replaced with the conditional *p53ER*^*TAM*^ allele ^67^. Since p53ER^*TAM*^ is functionally inactive in the absence of tamoxifen, *KPO* animals without tamoxifen are effectively p53^*null*^. This greatly accelerates spontaneous transition of KRas^*G12D*^-driven PanIN to PDAC ^17,18^. In addition, *KPO* animals harbor a doxycycline-inducible transgene encoding the dominant negative Myc dimerization mutant Omomyc that, upon induction, efficiently blocks endogenous Myc function ^68^. 10-week-old *KPO* mice harboring extensive, highly desmoplastic spontaneous PDAC were treated with doxycycline for 4 weeks to inhibit endogenous Myc. As with de-activation of MycER^T2^ in *KM*^*pdx*1^ and *KM*^*p*48^ mice, inhibition of endogenous Myc by Omomyc triggered quantitative regression of all spontaneous PDAC tumors together with their extensive attendant fibroinflammatory stroma (Figure 6B).

**Figure 6.**
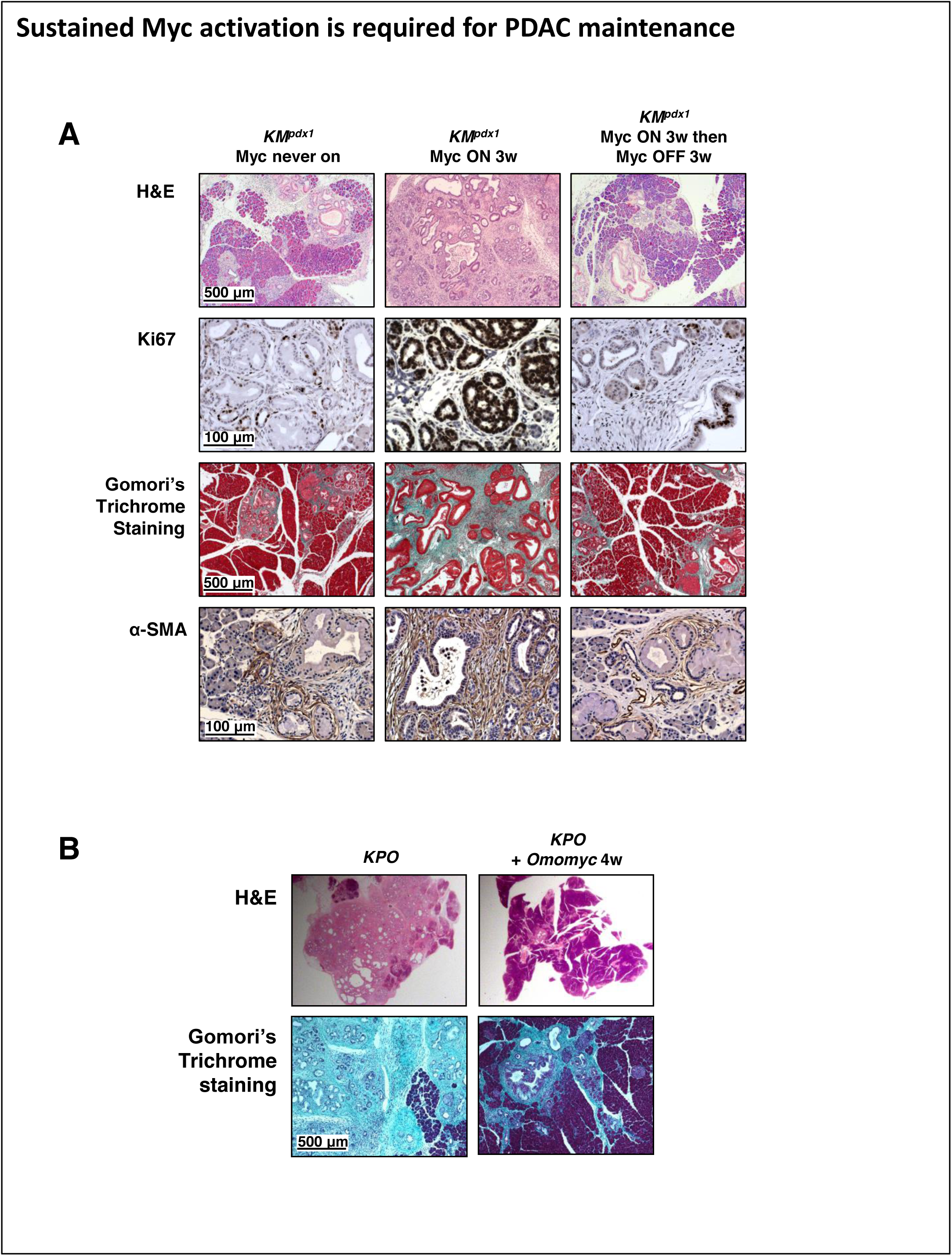
Sustained Myc activation is required for PDAC maintenance (related to Figure S7) (A) Representative H&E (top) and immunohistochemical staining for proliferation (Ki67, second row), Gomori’s Trichrome staining for collagen (third row) and α-SMA (bottom row) in sections of pancreata from 12 week old *KM*^*pdx*1^ mice in which Myc was never activated (treated with oil for 3 weeks and then left untreated for a further 3 weeks), or activated for 3 weeks (treated with tamoxifen for 3 weeks (middle), or activated for 3 weeks and then switched off (treated with tamoxifen for 3 weeks and then untreated for a further 3 weeks) (right). (B) Representative H&E (top) or Gomori’s Trichome stained (bottom) sections of pancreata from 10-week old *K*^*pdx*1^*; p53ER*^*TAM*^ *KI* (KPO) mice without (left) or with Omomyc expression for 4 weeks (right).

To define the detailed kinetics of PDAC regression following Myc de-activation, MycER^T2^ was activated for 3 weeks in 12-week-old *KM*^*pdx*1^ mice to generate aggressive pancreatic adenocarcinomas and then acutely de-activated by tamoxifen withdrawal for 1 and 3 days (Figure 7). Myc de-activation triggered immediate tumor cell arrest and progressive reversal of the PDAC signature stromal changes that Myc activity initially induced - specifically, rapid efflux of macrophages and neutrophils, rapid re-infiltration of CD3+ T cells, de-activation of stellate cells, dissolution of desmoplasia, re-emergence of patent vasculature and reversion from hypoxia back to normoxia (Figure 7). All of the above stromal reversion occurred with comparable rapidity to their initial Myc-driven formation and were accompanied by apoptosis of tumor cells and regression of tumors. Furthermore, regression with identical histology and kinetics was observed following Myc de-activation in PDAC-bearing *KM*^*p*48^ animals (Figure S8). In addition to the above straightforward reversals of various Myc-induced changes, however, PDAC regression also featured unique processes absent from either normal pancreas or PDAC. Most obviously, Myc de-activation triggered not only the proliferative arrest of proliferating tumor cells but also their apoptosis. Since analogous Myc deactivation in cultured *KM*^*pdx*1^ tumor cells hitherto maintained in the presence of 4-OHT triggered no such cell death (data not shown), it is evident that Myc deprivation is not inherently toxic to PDAC cells but rather a vulnerability manifest in tumor cells *in vivo*. PDAC regression was also marked by rapid and meticulous disassembly of the extensive juxta-PDAC desmoplasia and by the rapid migration of B220^+^ cells from the PDAC tumor periphery into the regressing tumor mass itself. To address whether PDAC regression was mediated by the influx of CD3^+^ T cells we systemically ablated CD4^+^ and CD8^+^ T cells. However, this had no impact on either rate or extent of PDAC regression (Figure 8). Hence, as recently shown in lung adenocarcinoma, CD4^+^/8^+^ T cells play no measurable role in PDAC regression.

**Figure 7.**
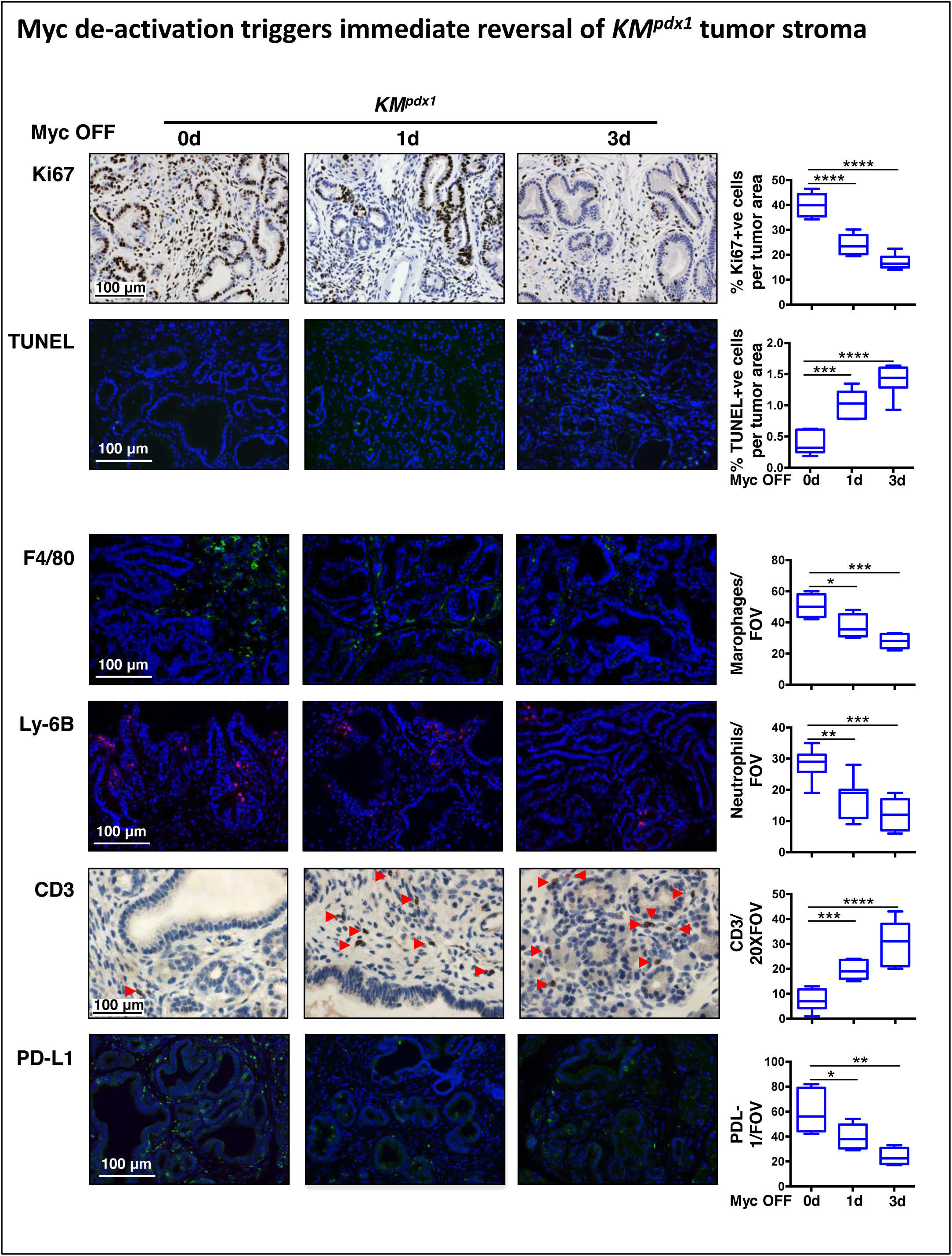
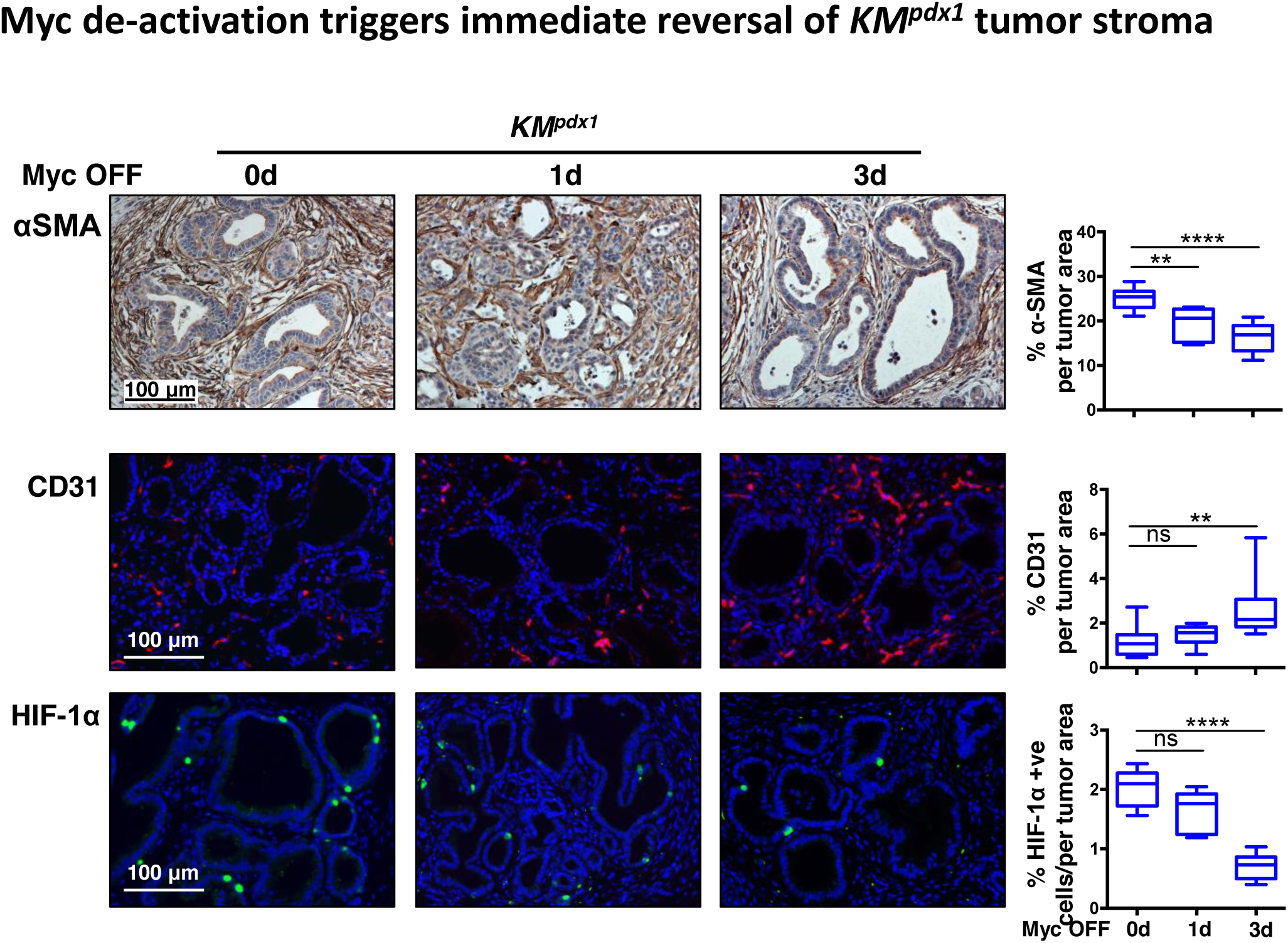
Myc de-activation triggers immediate reversal of *KM*^*pdx*1^ tumor stroma (related to Figure S8) Immunohistochemical staining (IHC) for proliferation (of Ki67), immunofluorescence (IF) staining of cell death (TUNEL), macrophages (F4/80) and neutrophils (Ly-6B), IHC of T cells (CD3), IF of PD-L1, IHC of α-SMA, and IF for endothelial cells (CD31) and hypoxia (HIF-1α) in sections of pancreata harvested from 12 week old *KM*^*pdx*1^ mice treated with tamoxifen (Myc ON) for 3 weeks (0 day, left) followed by tamoxifen withdrawal (Myc OFF) for 1 (middle) and 3 days (right). Scale bars apply across each row. Quantification is shown on the right. Data were analyzed using unpaired t-test. n = 4-6 for each treatment group. *p < 0.05, **p < 0.01, ***p < 0.001, ****p < 0.0001 (ns = non-significant). FOV = field of view.

**Figure 8.**
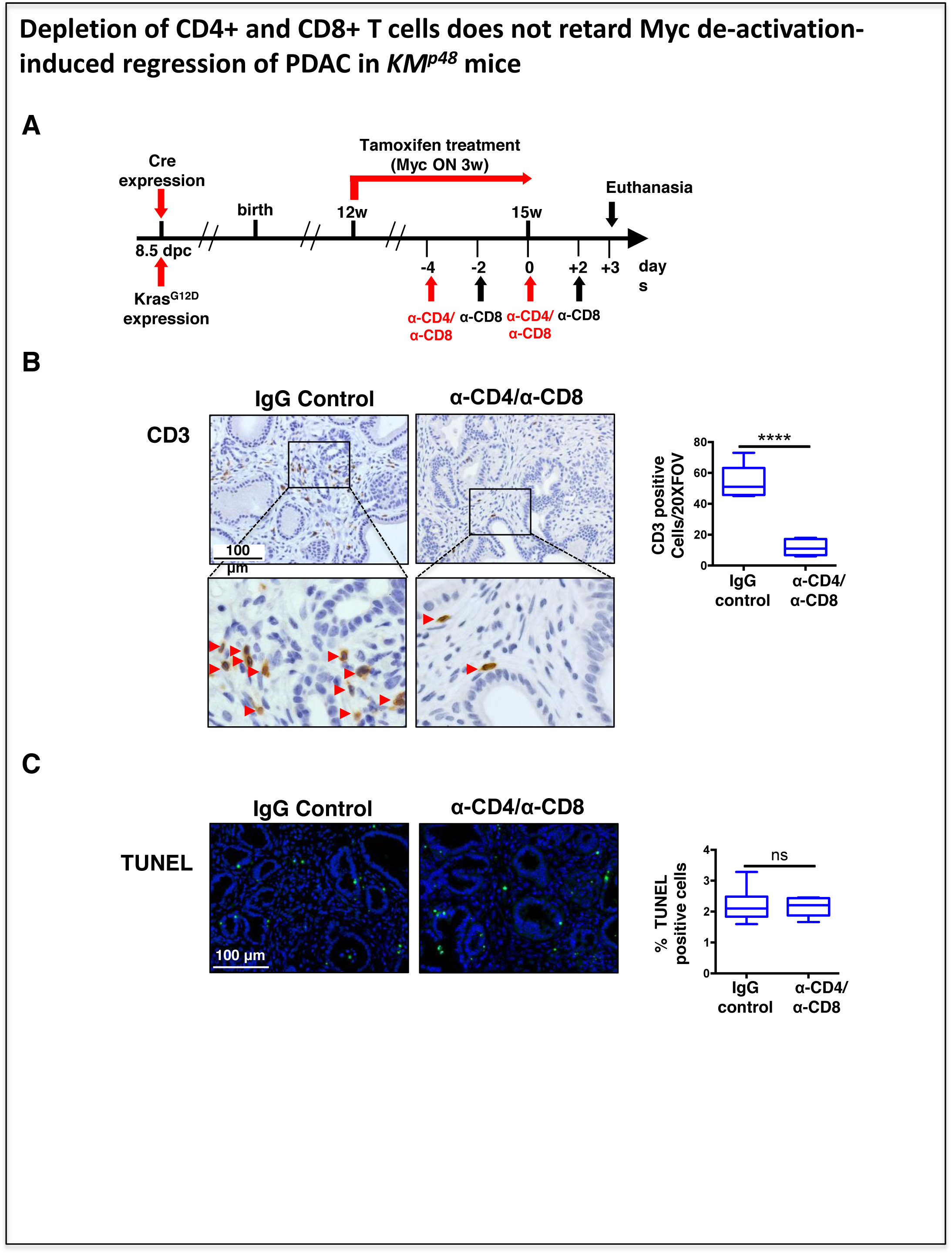
Depletion of CD4^+^ and CD8^+^ T cells does not retard Myc de-activation-induced regression of PDAC in *KM*^*p*48^ mice. (A) Schematic representation of CD4 and CD8 blocking experiment: 12-week old *KM*^*p*48^ mice were administered tamoxifen diet (Myc ON) for 3 weeks. Starting four days before cessation of tamoxifen diet (day 0, Myc OFF), mice were injected four times every other day (−4, −2, 0, +2 day time points) with neutralizing antibodies against CD8 (or IgG control) and injected twice four days apart (−4 and 0 day time points) with neutralizing antibodies against CD4 (or IgG control). Mice were euthanized three days post turning Myc off (+ 3 day). (B) Immunohistochemistry staining and quantification of CD3^+^T cells in pancreatic tumors isolated from *KM*^*p*48^ mice described in (a). (C) Immunofluorescence staining and quantification of cell death by TUNEL in in pancreatic tumors isolated from *KM*^*p*48^ mice described in (a). n = 4 per group. ****p < 0.0001 (ns = non-significant). Data were analyzed using unpaired t-test. FOV = field of view.

## DISCUSSION

Exhaustive studies indicate that activation of Myc is, alone, insufficient to drive transformation *in vitro* or oncogenesis *in vivo* and requires the cooperative input of other mitogenic mutations – notably activating mutations either in Ras itself or in the greater Ras signaling network ^25^. Oncogenic Ras, too, is a weak oncogene alone, typically generating indolent adenomatous lesions that only sporadically and rarely progress to adenocarcinoma through chance acquisition of additional oncogenic lesions. Obligate cooperative interdependence between Ras and Myc is recapitulated in pancreas *in vivo*. Cre recombinase-induced expression of KRas^*G12D*^ alone in pancreas progenitors drives the protracted and highly inefficient formation of indolent ductal lesions representing each of the three stages of PanIN. Activation of Myc alone in adult mouse pancreata has little discernible impact save for modest focal increases in proliferation. By contrast, activation of Myc in established KRas^*G12D*^-driven PanIN drives their uniform and immediate transition to aggressive pancreatic adenocarcinoma.

The tumor-stromal phenotype of the pancreatic adenocarcinomas induced by Myc activation exhibits several notable features. First, is the remarkable phenotypic resemblance of the Myc-induced adenocarcinoma phenotype to that of spontaneous PDAC, reproducing essentially all the canonical features of the most aggressive spontaneous human PDACs, including abundant infiltration of macrophages (F4/80^+^ and CD206^+^) ^69^, neutrophils and/or granulocytic MDSCs (Ly-6B^+^) ^59,70-72^ and B220^+^ B cells ^73-76^, absence of CD3+ T cells ^69^, activation (αSMA expression) and mitogenesis of stellate cells, extensive deposition of desmoplasia and concomitant blood vessel compression, vascular involution and tumor hypoxia ^8,77,78^. Second, is the complexity of the Myc-induced PDAC tumor-stromal phenotype, which involves the coordinated choreography of multiple, distinct stromal cell types - inflammatory, lymphoid, endothelial and mesenchymal cells – and commandeers almost all of the classical “hallmarks of cancer” ^79,80^. A third feature of the Myc induced PDAC phenotype is the immediacy of the stromal changes that Myc induces, most of which are clearly evident within 24 hours of MycER^T2^ activation. Such immediacy indicates that the PDAC phenotype is actively instructed and not a passive consequence of phenotypic constraints imposed by the surrounding host tissue. Since Myc in *KM*^*pdx*1^ and *KM*^*p*48^ mice is activated only in pancreas epithelial cells, we reasoned that the extensive stromal changes must be driven, and presaged, by instructive signals emanating from the those epithelial PanIN cells. Consistent with this, interrogation of pancreas extracts 6 hours after MycER^T2^ activation in *KM*^*p*48^ mice showed clear induction of a discrete ensemble of candidate signaling molecules. While our current analysis of signaling molecules is by no means as comprehensive as recent proteome-based studies ^81^, our kinetic data provide unique causal information as to both the sources and sequence of signals and responses that instigate the complex PDAC tumor-stromal conglomerate. One of the earliest Myc-induced stromal changes we observe is the rapid influx of F4/80 and CD206 macrophages consistent with the cotemporaneous induction of two well-described macrophage chemoattractants – CCL2 and CCL9. Macrophages have diverse origins and functions in normal pancreas ^82,83^ but in PDAC, where heavy macrophage infiltration is associated with poor prognosis, tumor-associated macrophages are known to promote EMT and tumor cell migration, suppress immunity and promote recruitment of myeloid-derived suppressor cells, drive myofibroblastic trans-differentiation of stellate cells and cancer-associated fibroblasts (CAFs) through release of multiple secondary signals including PDGF and TGFβ, and deployment of various proteases that actively remodel stroma ^55,83,84^. Inhibition of macrophages through blockade of the CCR2 receptor has been shown to enhance anti-tumor immunity, slow PDAC growth and reduce metastases ^55,71,85^. Myc activation in *KM*^*pdx*1^ and *KM*^*p*48^ PanIN also triggered the rapid influx of Ly6B^+^ neutrophils. Infiltration by neutrophils (and their G-MDSC kin) in PDAC is associated with the most undifferentiated, aggressive and metastatic forms of the disease with the poorest prognosis ^5:Felix, 2016 #349,86,87^ and is thought to be a consequence of their secretion of multiple stromal remodeling proteases and diverse secondary signaling molecules including IL-8, GM-CSF, CCL3, CXCL10, IL-2, TNFα and oncostatin-M ^59^, and through suppression of T cell surveillance ^70^. Several studies have indicated that a pivotal role is played by CXCL5 and its neutrophil receptor CXCR2 in recruitment of neutrophils in PDAC. Indeed, elevated expression of CXCL5 is associated with the most aggressive disease while blockade of either CXCL5 or CXCR2 impedes neutrophil accumulation in PDAC, slows PDAC growth ^70,71,86^ and promotes influx of T lymphocytes ^70^. The induction of CXCL5 (Figure 3) we observe within only 6 hours post Myc activation is entirely consistent with the subsequent neutrophil influx we observe during the Myc-driven transition to PDAC. However, prompted by the previously reported role of GAS6-AXL signaling in neutrophil dynamics ^88,89^ and the rapid Myc-dependent induction of GAS6 (Figure 3), a member of the Protein S family of vitamin K-dependent plasma glycoproteins, we uncovered an additional signaling pathway necessary for Myc-driven neutrophil influx. GAS6 is a ligand for the TAM receptor tyrosine kinase family (Tyro3, AXL and MER) and its induction by Myc temporally correlated with appearance of phospho-AXL in a proportion of PanIN epithelial cells. Blockade of either GAS6 with low dose Warfarin ^64,90^ or AXL kinase with the selective AXL kinase inhibitor TP-0903 ^66,91,92^ completely blocked Myc-induced neutrophil influx and markedly suppressed proliferation in both tumor cell and stromal compartments - most especially stellate cells - while exerting only a modest inhibitory impact macrophage infiltration (Figure 4). GAS6-AXL blockade also completely blocked the rapid influx of B220^+^ B cells to the peritumoral stroma that Myc triggers (Figure 4). A pivotal role for B lymphocytes has lately emerged in PDAC progression and aggression, most notably innate B1b cells, hitherto implicated in physiological processes such as microbial defense and removal of debris after tissue damage ^93,94^. B cells are attracted to incipient PDAC via CXCL13 and, through their release of IL-35, are thought to suppress T cell immunity ^95-97^, promote PDAC tumor cell proliferation and invasiveness and inhibit apoptosis ^98,99^. Diverse studies have demonstrated GAS6-AXL pathway activation in both tumor and stromal compartments of various solid tumors, including PDAC ^81^ and shown that GAS6-AXL pathway inhibition impedes growth of many cancers including PDAC ^100^ and promotes responses to chemotherapy ^65,91,92,100,101^. However, while GAS-AXL signaling is broadly implicated in regulating aspects of immunity ^61,88,92,102-104^ and tumor-stromal signaling ^81^, there is no previously described involvement of GAS6-AXL signaling in either neutrophil or B cell dynamics in PDAC. Presently, it remains unclear what the relative contribution of inhibition of each of the individual target cell populations of GAS6-AXL blockade - neutrophils, B lymphocytes or stellate cells – makes to Myc-driven PDAC progression, or whether each of these targets is affected independently or some are contingent upon another. Concomitantly with the influx of macrophage, neutrophils and B cells, Myc activation induced immediate expulsion of CD3+ T lymphocytes from PanINs and this coincided with rapid induction of surface PD-L1 on PanIN epithelial cells *in vivo*. PD-L1 was also induced by Myc in isolated *KM*^*pdx*1^ tumor cells *in vitro*, where ChIP analysis located Myc on the *CD274* (*PD-L1*) gene promoter. The immediacy and cell autonomy of Myc-induced PD-L1 induction in PanIN epithelium both intimate that *CD274* gene is a direct Myc target in pancreas, as suggested in some other tissues ^105^. Systemic blockade of PD-L1 with an appropriate antibody completely abrogated Myc-induced T cell efflux, demonstrating PD-L1 to be causally required for T cell efflux. However, the consequent persistence of intra and peritumoral CD3+ T cells exerted no discernible inhibitory impact on either PDAC cell survival or PDAC growth (not shown). These data are in accord with previous studies in both pancreas ^106^ and lung ^54^ that indicate that enforced persistence of T cells in tumors is, alone, insufficient to engage significant anti-tumor immune responses. In spontaneous PDAC, activated stellate cells are known to be the principal agents of desmoplastic deposition ^8,56,57,81,107^. While the role of PDAC desmoplasia in vascular compression and hypoxia is generally accepted ^8,9,11,56,58,77,78,108^, its influence on disease progression and response to therapy remain contentious ^11,109-112^. Upon acute activation of Myc, peritumoral *KM*^*pdx*1^ and *KM*^*p*48^ pancreatic stellate cells undergo trans-differentiation into highly proliferative, αSMA^+^ myofibroblasts. This is accompanied by progressive desmoplasia, vascular compression and onset of hypoxia and demonstrates that desmoplastic deposition is hard-wired into PDAC progression.

The final notable feature of the Myc-driven PDAC phenotype is its remarkable tissue specificity, which is quite distinct from that of adenocarcinomas induced in lung by the exact same combination of KRas^*G12D*^ and switchable Myc {Kortlever, 2017 #312} (Table 1). In both pancreas and lung, the Myc-induced adenocarcinoma phenotype arises immediately following Myc activation, involves rapid onset of tumor cell proliferation and invasion, influx of macrophages and T cell exclusion, and closely mirrors the phenotypes of spontaneous adenocarcinomas arising each tissue. However, the two phenotypes differ dramatically in other respects, a difference clearly reflected in the signaling molecules deployed in each instance. Thus, rapid influx of macrophages is common to the Myc-induced adenocarcinoma transition in both pancreas and lung, consistent with the pivotal role such cells play in tissue remodeling, and this is reflected in significant commonality in the induction of the monocyte attractant CCL9, augmented in pancreas by co-induction of CCL2. Rapid, Myc-induced CD3^+^ T cell exclusion is also seen in both tissues and, in both cases, is dependent upon induction of PD-L1. However, the mechanism of PD-L1 deployment is tissue-specific. In pancreas, Myc induces expression of PD-L1 cell autonomously on Myc-expressing PanIN epithelial cells, most likely through direct Myc-dependent transcription. By contrast, PD-L1 is never appreciably induced in Myc-expressing lung adenoma epithelial cells themselves and is, instead, ported into incipient tumors on incoming alveolar macrophages. Another example of tissue-specificity in response to Myc is the dramatic and immediate influx of neutrophils induced by Myc in PanINs, together with activation of its requisite GAS6-AXL pathway. Both neutrophil influx and pAXL are entirely absent from lung (Figure 4 and data not shown), a tissue with already high basal levels of neutrophils. NK and B220+ B cells also exhibit different dynamics following Myc activation in each tissue: peritumoral NK cells are rapidly expelled from lung adenomas but are essentially absent from pancreas tumors while Myc induces B cell expulsion from lung adenomas but rapid influx of B cells to the tumor periphery in PanINs. Perhaps most dramatically, Myc activation in PanIN induces rapid activation and proliferation of local fibroblastic and stellate cell in pancreas, together with abundant deposition of desmoplastic stroma, blood vessel compression and hypoxia. By contrast, Myc activation in lung elicits no appreciable activation of local fibroblastic cells or significant desmoplasia and a dramatic pro-angiogenic switch fueled by macrophage-derived VEGF, which rapidly restores normoxia to the otherwise indolent, hypoxic adenomas ^54^.

The remarkable tissue-specificity of the complex, multi-cell lineage tumor-stroma phenotypes that the combination of KRas^*G12D*^ and Myc instruct, together with the remarkable similarity of those phenotypes to those of spontaneous tumors arising in each of the same tissues, strongly supports the idea that adenocarcinoma phenotypes are principally an attribute of their host tissue rather than their peculiar ensemble of driving oncogenes. In this scenario, Ras and Myc serve as common downstream conduits of the multitude of different oncogenic mutations that lie upstream and, rather than specifying the tumor phenotype, KRas^*G12D*^ and Myc instead unlock a preconfigured tissue-resident program. This notion is entirely consistent with recent studies showing that tumor cell-intrinsic factors dictate aspects of stromal biology, most notably immune dynamics ^113^. The likely source of this tissue-resident program is suggested by the remarkable phenotypic overlap between PDAC (spontaneous and Myc-driven) and that of regenerating pancreas following injury. Both exhibit marked acinar-ductal metaplasia, epithelial proliferation, rapid infiltration of macrophages and neutrophils, exclusion of T cells, activation of stellate cells and an extreme fibroinflammatory reaction ^114-117^, attributes so overtly similar as to frequently confound differential diagnosis between pancreatitis and early PDAC. Hence, we conclude that oncogenic Myc “hacks” the endogenous pancreas regenerative program.

The rapid regression of advanced, invasive *KM*^*pdx*1^ and *KM*^*p*48^ PDACs upon Myc de-activation, and of spontaneous KRas^*G12D*^-induced (and p53-deficient) mouse PDAC upon inhibition of endogenous Myc, indicates that the PDAC phenotype-notwithstanding its complexity, multifarious component cell types and abundant desmoplasia - is inherently reversible. Indeed, sustained blockade of Myc function induces the quantitative regression of even the most extensive tumor masses, restoring normal exocrine pancreas microarchitecture, albeit with a macroscopic organ structure badly disfigured by scars and cysts. There are several features of note in this remarkable PDAC regression. First, the onset of regression is immediate upon MycER^T2^ deactivation, proceeding with rapid kinetics comparable to those with which Myc initially drove PDAC progression. Such rapidity indicates that, like Myc protein and mRNA, key cell intrinsic and extrinsic pro-tumorigenic effectors downstream of Myc are similarly ephemeral, attenuating rapidly unless continuously re-engaged by Myc activity. Second, since Myc is de-activated only in the PDAC epithelial compartment, all induced stromal alterations associated with regression must be initiated by altered signaling originating in the PDAC epithelium. Moreover, we see no evidence whatsoever that any aspect of the PDAC tumor stroma is independent of Myc and self-sustaining. Third, macroscopic tumor regression, like its Myc-driven progression counterpart, proceeds via a complex choreography across multiple diverse cell types that, in the main, walks back the processes Myc initially induced: tumor cells withdraw from cycle and resume their prior acinar disposition, macrophages and neutrophils rapidly exit, PD-L1 expression falls and CD3+ T cells re-infiltrate, stellate cells de-activate and arrest. After a brief lag, the consequent progressive disassembly of desmoplasia relieves interstitial pressure, restoring patency to blood vessels and tissue reoxygenation. Nonetheless, regression is not a simple reversal of Myc-driven progression since it invokes processes absent from both normal pancreas and PDAC. Most obviously, Myc de-activation triggers not only arrest of proliferating tumor cells but also their death. PDAC regression also requires meticulous disassembly of the extensive PDAC desmoplasia and is accompanied by unexpected migration of B220+ cells from the PDAC tumor periphery into the regressing tumor mass itself. As with Myc-driven PDAC genesis, the rapidity, reproducibility, efficiency and tight choreography of PDAC regression all suggest it is a manifestation of some coordinated physiological process. Clues to the origin of this process come from multiple studies of the post-regenerative resolution phase of pancreas injury by which normal pancreas mass, architecture and function are restored. Such injury resolution involves ductal to acinar transdifferentiation and the involution of fibrosis, and requires both the rapid exclusion of F4/80 macrophages and neutrophils and influx of lymphocytes ^114,121,122^. It seems plausible that, just as activating Myc hacks into the endogenous pancreas regenerative program, inhibiting Myc (or upstream oncogenic signals) hacks into a physiological injury resolution program that evolved to terminate post-injury regeneration, prune excess cells and reinstate normal tissue architecture. Indeed, it is possible that enforced engagement of this same injury resolution program is the principal mechanism by which targeted inhibition of oncogenic signaling exerts its curative impact in cancers.

## MATERIALS AND METHODS

### Generation and maintenance of genetically engineered mice

*LSL-Kras*^*G12D*^, *p48*^*cre/+*^, *Pdx-1-Cre, p53ER*^*TAM*^, *TRE-Omomyc*, and *Rosa26-lsl-MER*^*T*2^ mice have all been previously described (Jackson 2001, Kawaguchi 2002, Hingorani 2003, Christophorou 2005, Soucek, 2008; Murphy 2008). *CMVrtTA* [*Tg(rtTAhCMV)4Bjd/J*] mice were purchased from Jackson Laboratories. Mice were maintained on a 12-hour light/dark cycle with continuous access to food and water and in compliance with protocols approved by the UK Home Office guidelines under a project license to G.I.E. at the University of Cambridge. Deregulated Myc activity was engaged in pancreatic epithelia of *KM*^*pdx*1^ and *KM*^*p*48^ mice by daily intraperitoneal (i.p.) administration of tamoxifen (Sigma-Aldrich, TS648) dissolved in sunflower oil at a dose of 1 mg/20 g body mass; sunflower oil carrier was administered to control mice. *In vivo*, tamoxifen is metabolized by the liver into its active ligand 4-Hydroxytamoxifen (4-OHT). For long-term treatment, mice were placed on tamoxifen diet (Harlan Laboratories UK, TAM400 diet) as described ^118^; control mice were maintained on regular diet. Omomyc expression was systemically induced in *KPO* (*pdx1-Cre;LSL-KRas*^*G12D*^*; p53ER*^*TAM*^ *KI;TRE-Omomyc;CMVrtTA*) mice by addition of doxycycline (2 mg/ml plus 5% sucrose) to their drinking water versus 5% sucrose for control mice.

### Tissue preparation and histology

Mice were euthanized by cervical dislocation and cardiac perfused with PBS followed by 10% neutral-buffered formalin (Sigma-Aldrich, 501320). Pancreata were removed, fixed overnight in 10% neutral-buffered formalin, stored in 70% ethanol and processed for paraffin embedding. Tissue sections (4 μM) were stained with hematoxylin and eosin (H&E) using standard reagents and protocols or with Gomori’s Trichrome Stain (Polysciences, 24205), according to the manufacturer’s instructions. For frozen sections, pancreata were embedded in OCT (VWR Chemicals, 361603E), frozen on dry ice and stored at –80°C.

### Immunohistochemistry and immunofluorescence

For immunohistochemical (IHC) or immunofluorescence (IF) analysis, paraffin-embedded sections were de-paraffinized, rehydrated, and either boiled in 10 mM citrate buffer (pH 6.0) or treated with 20ug/ml Proteinase K to retrieve antigens, depending on the primary antibody used. For IF studies, OCT-embedded sections were air-dried and fixed for 30 min in 1% paraformaldehyde. Primary antibodies used were as follows: rabbit monoclonal anti-Ki67 (clone SP6) (Lab Vision, Fisher, 12603707), rat anti-ki67 (Clone SolA15, Fisher, 15237437), rat monoclonal anti-CD45 (clone 30-F11, BD Pharmingen, 553076), rat monoclonal anti-neutrophils (Clone 7/4, Cedarlane, CL8993AP), rat monoclonal F4/80 (clone Cl:A3-1, Bio-Rad, MCA497R); goat polyclonal anti-MMR/CD206 (R&D Systems, AF2535); rabbit monoclonal anti-CD3 (clone SP7, ThermoFisher, RM-9107-RQ), rabbit polyclonal anti-CD274/PD-L1 (Abcam, ab58810); rat monoclonal anti-CD45R/B220 (clone RA3-6B2, ThermoFisher, MA1-70098), rabbit polyclonal anti-αSMA (Abcam, ab5694), rabbit polyclonal CD31 (Abcam, 28364), rabbit polyclonal anti-Hif1-α (Abcam, ab82832), rabbit polyclonal anti-pAXL (Y779, AF2228 R&D Systems), mouse monoclonal anti Pan-Keratin (clone C11, NEB, 4545S). Primary antibodies were incubated with sections overnight at 4°C except for anti-CD3, which was applied for 20 minutes at room temperature. For IHC analysis, primary antibodies were detected using Vectastain Elite ABC HRP Kits (Vector Laboratories: Peroxidase Rabbit IgG PK-6101, Peroxidase Rat IgG PK-6104, Peroxidase Goat IgG PK-6105) and DAB substrate (Vector Laboratories, SK-4100); slides were then counterstained with hematoxylin solution (Sigma-Aldrich, GHS232). For IF analysis, primary antibodies were visualized using species-appropriate cross-adsorbed secondary antibody Alexa fluor 488 or 555 conjugates (ThermoFisher scientific); slides were counterstained with Hoechst (Sigma-Aldrich, B2883) and mounted in Prolong™ Gold Antifade Mountant (ThermoFisher scientific; P36934). TUNEL staining for apoptosis was performed using the Apoptag Fluorescein *in situ* Apoptosis Detection Kit (Millipore; S7110) according to manufacturer’s instructions. Images were collected with a Zeiss Axio Imager M2 microscope equipped with Axiovision Rel 4.8 software.

### Mouse inflammation/Cytokine antibody Arrays and quantitative real time-PCR

Pancreata were collected at appropriate time points and snap-frozen in liquid nitrogen. Whole pancreas protein extracts were isolated and incubated with mouse inflammation/cytokine antibody arrays (40 targets, Abcam, ab133999; 97 targets, Abcam ab169820), according to manufacturer’s instructions. Signal intensity was determined using Image J software. Differential expression of cytokine RNA was assessed by quantitative real time-PCR (RT-PCR). For this, total pancreas RNA was isolated using a Qiagen RNeasy Plus Isolation Kit followed by cDNA synthesis (High Capacity cDNA RT kit, Applied Biosystems, 4374966). RT-PCR was performed using TaqMan Universal Master Mix II (Fisher, 4440038), according to manufacturer’s protocol. Primers used were: *Cxcl5* (Fisher, Mm00441260_m1), *Gas6* (Fisher Mm00441242_m1), *Ccl9* (Fisher, Mm00436451_g1), *Ccl2* (Fisher, Mm00490378_m1), *Tbp* (Fisher, Mm00446973_m1), *Shh* (Fisher, Mm00436528_m1). Samples were analyzed in triplicate on an Eppendorf Mastercycler Realpex 2 with accompanying software. *Tbp* was used as an internal amplification control.

### Isolation of primary mouse pancreatic cell lines and chromatin Immunoprecipitation (ChIP)-qPCR analysis

Pancreatic tumors were isolated as previously described (Olive 2009), but with modifications. Briefly, 12 week-old *KM*^*pdx*1^ and *KM*^*p*48^ mice were placed on tamoxifen diet for 3 weeks. Pancreatic adenocarcinomas were then harvested, minced, and digested at 37°C in Hank’s balanced salt solution containing 2 mg/ml type V collagenase (Sigma-Aldrich, C9263) for 30 min under constant agitation. Digested tumors were centrifuged and the pellet further digested with 1X Trypsin-EDTA (0.05% trypsin, 0.02% EDTA) (Sigma-Aldrich, 59418C) for 5 min. Proteases were inactivated by the addition of Dulbecco’s modified Eagle medium (DMEM) (ThermoFisher Scientific, 41966-029) containing 10% fetal bovine serum (ThermoFisher Scientific, 10270106). Established cell lines were propagated in DMEM containing 10% FBS and maintained with 100nM of (Z)-4-hydroxytamoxifen (4-OHT) (Sigma-Aldrich H7904) and routinely evaluated for mycoplasma contamination. For gene expression analysis, total RNA was isolated, reverse transcribed and amplified by RT-PCR as described above. Primers used were for *Cd274/pdl1* (Fisher, Mm00452054_m1), *Tbp* (Fisher, Mm00446973_m1). *Tbp* was used as normalization control.

ChIP was performed as described in (Carey, 2009), with modifications. Briefly, 10^7^ cells cultured with 100nM 4-OHT or ethanol control for 6 or 24 hours were incubated with 1% formaldehyde (Sigma-Aldrich, F8775) at room temperature for 10 min. Fixation was quenched by addition of glycine to a final concentration of 125mM for 5 min at room temperature. Cells were then washed with ice-cold PBS, pelleted and resuspended in lysis buffer (10mM Tris-HCl pH8.0, 85mM KCl, 0.5% NP40, 1mM DTT) with protease inhibitors (Sigma-Aldrich, 11836153001). Cross-linked lysates were sonicated to shear DNA to an average fragment size of 100 to 500bp. The following antibodies were then used for immunoprecipitation: rabbit polyclonal anti-c-Myc (CST, 9402), rabbit polyclonal anti ERα (clone HC20) (Santa Cruz, sc-543), normal rabbit IgG as a background control (CST, 2729). Chromatin regions enriched by ChIP were then identified by qPCR in triplicate using SYBR Green Reaction Mix (ThermoFisher Scientific, 4385616) according to the manufacturer’s protocol. *Pdl1* primers used for amplification were 5’-GTTTCACAGACAGCGGAGGT-3’ (forward) and 5’-CTTTAAAGTGCCCTGCAAGC-3’ (reverse) with NCBI reference sequence NM_021893.3 and described in (Noman, JEM 2014). Calculations of average cycle threshold (Ct) and standard deviation for triplicate reactions were performed and each DNA fraction normalized to its input to account for chromatin sample preparation differences.

### Blocking antibodies, drug preparation, and drug study treatment groups

For PD-L1 blocking studies, *KM*^*p*48^ mice were injected i.p. with 160 µg of LEAF-purified anti-mouse CD274/PD-L1 (clone 10F.9G2, Biolegend, 124309) or Rat IgG2b isotype control (Clone RTK4530, Biolegend, 400644) every 2 days for 2 weeks, starting one day prior to Myc activation. Mice were euthanized after 2 weeks and pancreata harvested for histology. For CD4/CD8 blocking studies, *KM*^*p*48^ mice were administered tamoxifen diet for 3 weeks. Commencing four days prior to returning the mice to normal (tamoxifen-free) diet they were injected twice i.p. with 200ug of rat anti-mouse CD4 (Clone GK1.5, Bio X Cell, BE003-1) four days apart, and four times with 200ug rat anti-mouse CD8 (Clone 2.43, Bio X Cell, BE0061) every other day. Control mice were injected with an equivalent amount of rat IgG2b isotype control (Clone LTF-2, Bio X Cell, BE0090). Mice were euthanized 3 days post Myc inactivation and pancreata harvested for histological analysis.

For Warfarin blockade of Gas6, *KM*^*p*48^ mice were randomized to receive either normal drinking water or water supplemented with 1 mg/L warfarin (Sigma-Aldrich, A2250) starting one day prior to Myc activation. Warfarin-containing water was replenished every 2 days. Mice were euthanized 3 days following Myc activation and pancreata harvested for histology. For AXL receptor tyrosine kinase inhibition, the specific inhibitor TP-0903.tartrate (Tolero Pharmaceuticals, Inc.) was dissolved in its formulation 5% (w/v) Vitamin E TPGS + 1% (v/v) Tween 80 in water and 55 mg/kg administered by oral gavage to *KM*^*p*48^ mice every 2 days, starting 2 hours before initial Myc activation. Mice were euthanized 5 days later and pancreata harvested for histological analysis.

### ***In vivo* ultrasound imaging**

*KM*^*p*48^ mice were anaesthetized by continuous infusion of 3% isofluorane using a veterinary anesthesia system (VetEquip, California). The ventral, lateral and dorsal abdominal surfaces were shaved and mice then injected i.p. with 2 ml of sterile saline. Tumors were imaged using a Vevo 2100 imaging system (Visual Sonics, Toronto, Canada). Serial images were collected at 0.2 mm intervals throughout the entire tumor. Vevo2100 v1.6.0 software was used to assess tumor size at each time point from 3D scans, identifying in the Z plane and calculating maximal width and length.

### Quantification and Statistical Analysis of tumor burden

For quantification of tumor burden, H&E sections were scanned with an Aperio AT2 microscope (Leica Biosystems) at 20X magnification (resolution 0.5 microns per pixel) and analyzed with Aperio Software. Statistical significance was assessed by Mann-Whitney U test, with the mean values and the standard error of mean (SEM) calculated for each group using Prism GraphPad software. Kaplan-Meier survival curves were calculated using the survival time for each mouse from each group, with the log-rank test used to calculate significant differences between the groups using Prism GraphPad software. Student’s t-test was employed in statistical analyses of ChIP and RT-PCR. IHC and IF staining were quantified using Fiji open source software; statistical significance was determined by Student’s t-test using Prism GraphPad software; data were represented as Box-and whisker plots, which show upper extreme, upper quartile, median, lower quartile, lower extreme. p values (ns = non-significant; *p < 0.05, **p < 0.01, ***p < 0.001, ****p < 0.0001) for each group (the number of animals in each cohort varied between 4-6 for each category).

## Author contributions

N.M.S and G.I.E. conceived the project. N.M.S, R.M.K., L.B.S., L.S., T.D.L., and G.I.E. designed experiments. N.M.S performed all experiments. R.M.K., V.J.A.B., L.P., T.C. and S.K. assisted with some experiments. M.J.A. did the histological analysis. N.M.S, T.D.L., and G.I.E. wrote the manuscript. G.I.E. supervised the study. All authors discussed results and revised the manuscript.

## Acknowledgments

We thank the members of the Evan laboratory, Drs Daniel Van Hoff (TGEN), Ronald Evans and Michael Downes (Salk Institute), Aarthi Gopinathan and Christine Feig (Cambridge Cancer Institute), Daniel Masso (Vall d’Hebron Institute), and Fernanda Kyle Cezar (King’s College London) for invaluable discussion and advice. We thank Alessandra Perfetto and Daniel Masso for assistance with animals, Stephanie Whike for assistance with histology and the Pre-clinical Genome Editing core at the Cambridge Cancer Research Institute (CI). We thank Tolero Pharmaceuticals, Inc. for their generous gift of TP-0903. The study was supported by program grants to G.I.E. (Cancer Research UK C4750/A12077 and C4750/A19013A, European Research Council (294851), and the SU2C/CRUK/Lustgarten Foundation Transatlantic Pancreatic Cancer Dream Team C4750/A22585).

