## Supplemental figures and table for "Myc instructs and maintains pancreatic adenocarcinoma phenotype"

Click here to download Supplemental Text and Figures Sodir et al. Supplementary Figures and text.pdf

#### Supplementary Figure Legends

##### Figure S1 (related to Figure 1)

**Myc drives progression to pancreatic adenocarcinoma in the *KM*$^{p48}$ mouse PDAC model** Representative H&E stained sections (top row), Immunohistochemical staining (IHC) for Ki67 (second row), α-SMA (third row) and Gomori's Trichrome staining (blue-stained collagen, fourth row) and CD45 immunofluorescence (IF) (red, bottom row) of sections of pancreata harvested from 15 week old *KM*$^{p48}$mice treated with either oil (Myc OFF, left) or tamoxifen (tam) (Myc ON, middle) and wild type control mice treated with tamoxifen (No Myc, right) for 3 weeks. Scale bars apply across each row.

### Myc drives progression to pancreatic adenocarcinoma in the *KM*$^{p48}$ mouse PDAC model

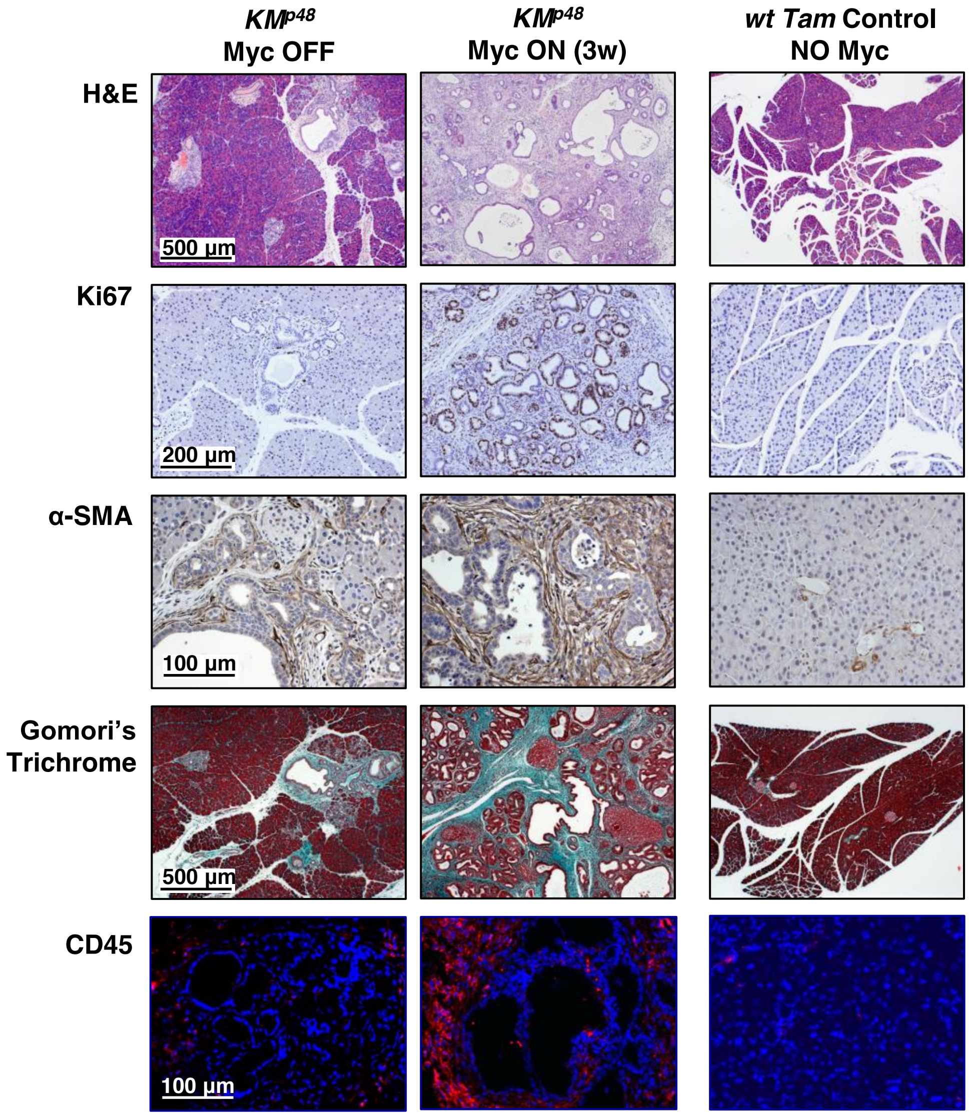

**Figure S2 (related to Figure 2)**

**Myc instructs immediate transition to pancreatic adenocarcinoma in *KM*$^{pdx1}$ mice**

Immunohistochemical staining (IHC) for proliferation (Ki67), immunofluorescence (IF) staining of macrophages (F4/80) and neutrophils (Ly-6B), IHC of T cells (CD3), and α-SMA, Gomori's Trichrome staining of collagen, and IF for endothelial cells (CD31) and hypoxia (HIF-1α) in sections of pancreata harvested from 12 week old *KM*$^{pdx1}$ mice treated with oil (Myc OFF, left panel) or tamoxifen for 1 (Myc ON, middle), 3 days (Myc ON, middle) or 7 days (Myc ON, right). Data were analyzed using unpaired t- test. Scale bars apply across each row. Quantification (Box-and whisker plots; right) shows upper extreme, upper quartile, median, lower quartile, lower extreme. n = 4-6 for each treatment group. $*p < 0.05$, $**p < 0.01$, $***p < 0.001$, $****p < 0.0001$ (ns = non-significant). FOV = field of view.

Sodir *et al*. Figure S2 (related to Figure 2)

#### Myc instructs immediate transition to pancreatic adenocarcinoma in *KM^pdx1* mice

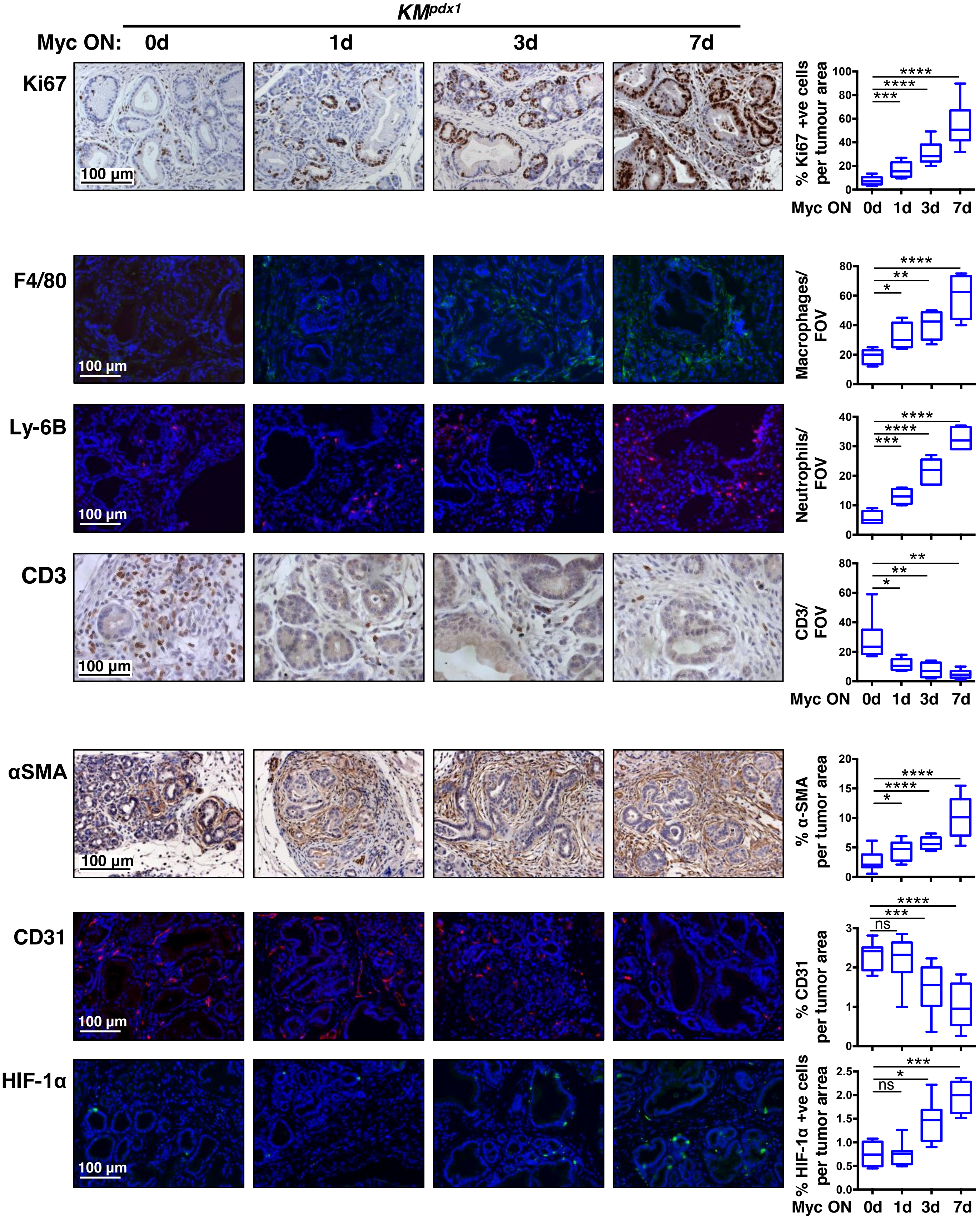

**Figure S3 (related to Figure 1)**

**Activation of Myc induces widespread intestinal neoplasia in *KM*$^{pdx1}$ mice causing premature death**

(A) Extra-pancreatic cooperation of Myc and *KRas*$^{G12D/+}$ in the small intestine of *KM*$^{pdx1}$ mice elicits rapid morphological changes and promotes intestinal neoplasia. Representative macroscopic images of 13-week old wild type tamoxifen treated control mouse (No Myc, left) and *KM*$^{pdx1}$ mice treated with either oil (Myc OFF, middle) or tamoxifen for 1 week (Myc ON, right). The red arrow indicates the duodenal swelling.

(B) H&E stained duodenal sections of 13-week old *KM*$^{pdx1}$ mouse treated with oil demonstrating normal (i), mild hyperplasia (ii) or aberrant monocryptal adenomas (iii). Activation of Myc for one week shows altered crypt-villus architecture (iv) (crypts are elongated and densely packed; villi are wider with branching, with markedly reduced luminal spaces in between and assuming a less differentiated morphology) and promoted tumorigenesis with neoplasia and hyperplasia shown in boxed areas with higher magnification (v) and (vi).

#### Activation of Myc induces widespread intestinal neoplasia in *KM*$^{pdx1}$ mice causing premature death

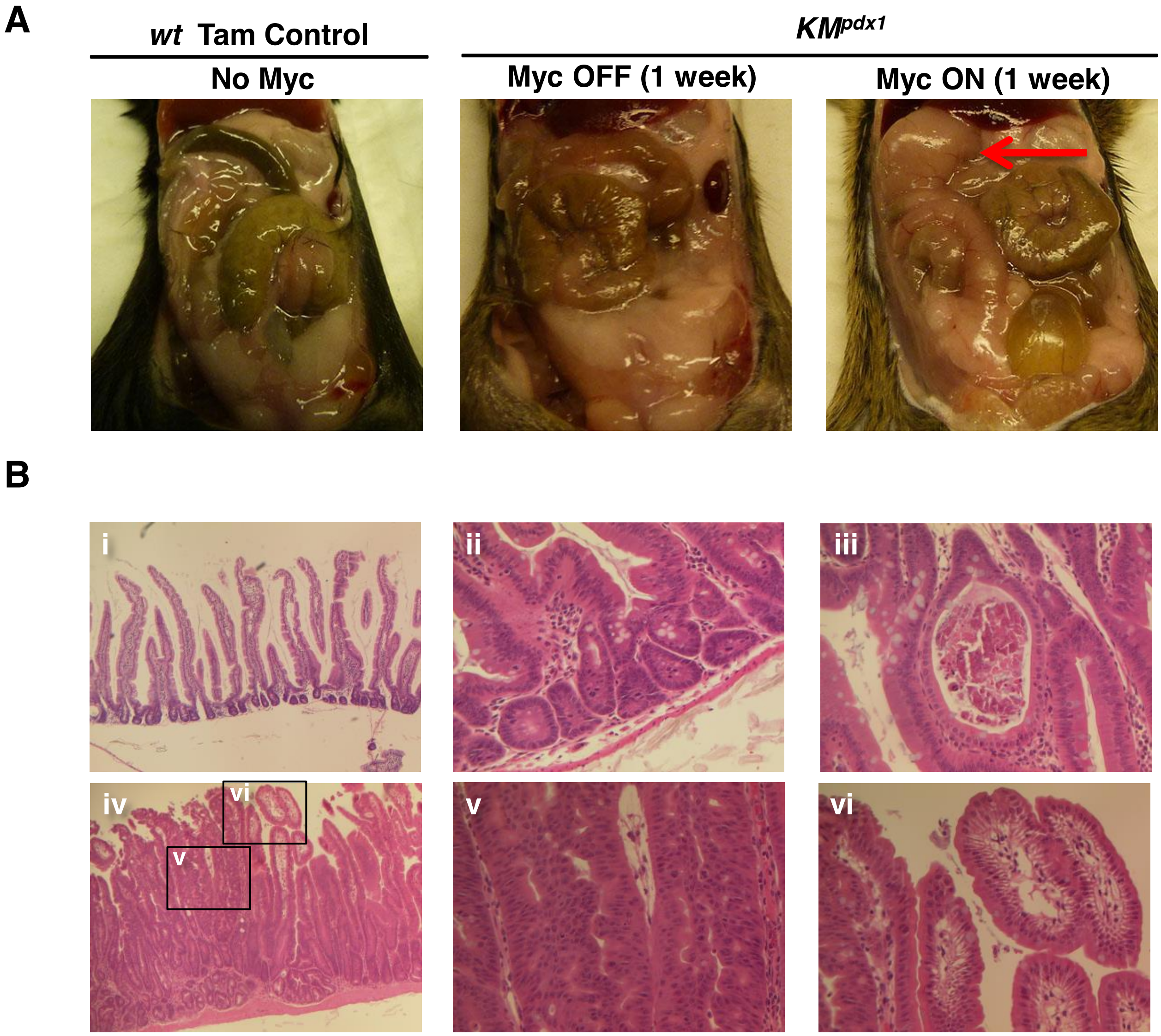

**Figure S4 (related to Figure 1)**

**Sustained Myc activation in the pancreas of *KM*$^{p48}$ mice induces lethal PDAC**

Kaplan-Meier survival curve (left) for *KM*$^{p48}$ cohorts either untreated (n=11, blue line) or treated with tamoxifen (tam) starting from 12 weeks of age (n=10, red line). Median survival with tamoxifen = 4.5 weeks. Log-rank (Mantel-Cox) test $p<0.0001$. The red arrow indicates a *KM*$^{p48}$ mouse whose pancreas is shown in the image to the right with a representative H&E stained section showing a PDAC. Boxed region is shown at higher magnification below.

Sodir *et al.* Figure S4 (related to Figure 1)

#### Sustained Myc activation in the pancreas of *KM*$^{p48}$ mice induces lethal PDAC

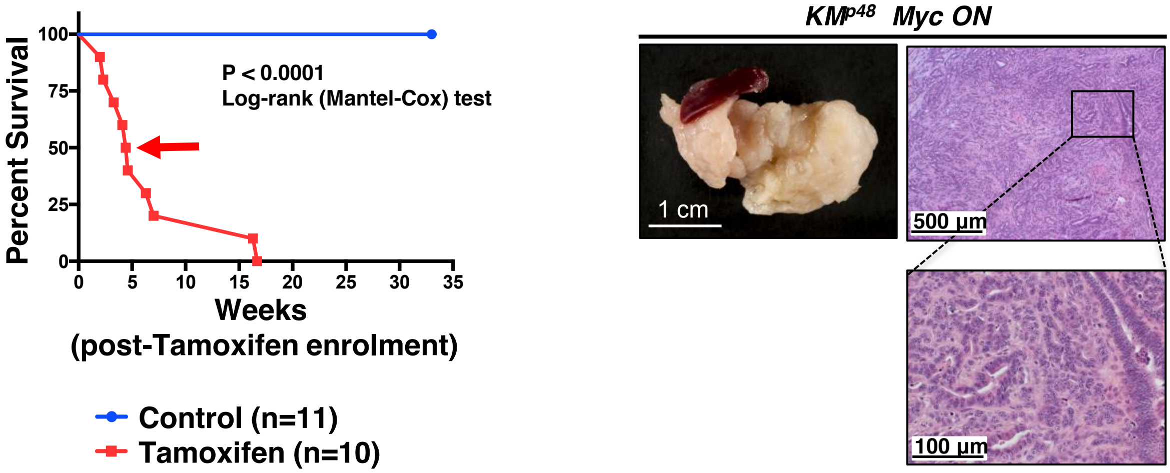

**Figure S5 (related to Figure 4)**

**Differential impact on various stromal compartments of inhibiting GAS6/AXL signaling in *KM*$^{p48}$ pancreatic tumors**

Representative double immunofluorescence staining of proliferating neutrophils (Ki67/Ly-6B), macrophages (Ki67/CD206) and $\alpha$-SMA (Ki67/$\alpha$-SMA), harvested from 12-week old *KM*$^{p48}$ mice untreated (left) or treated (right) with pAXL inhibitor TP-0903 concurrent with tamoxifen administration (Myc ON). Scale bar on bottom left applies to all large panels.

Quantification is shown on the right. n = 4-6 per group. ***$p < 0.001$, ****$p < 0.0001$. FOV = field of view.

Sodir *et al*. Figure S5 (related to Figure 4)

#### Differential impact on various stromal compartments of inhibiting GAS6/AXL signaling in *KM*$^{p48}$ pancreatic tumors

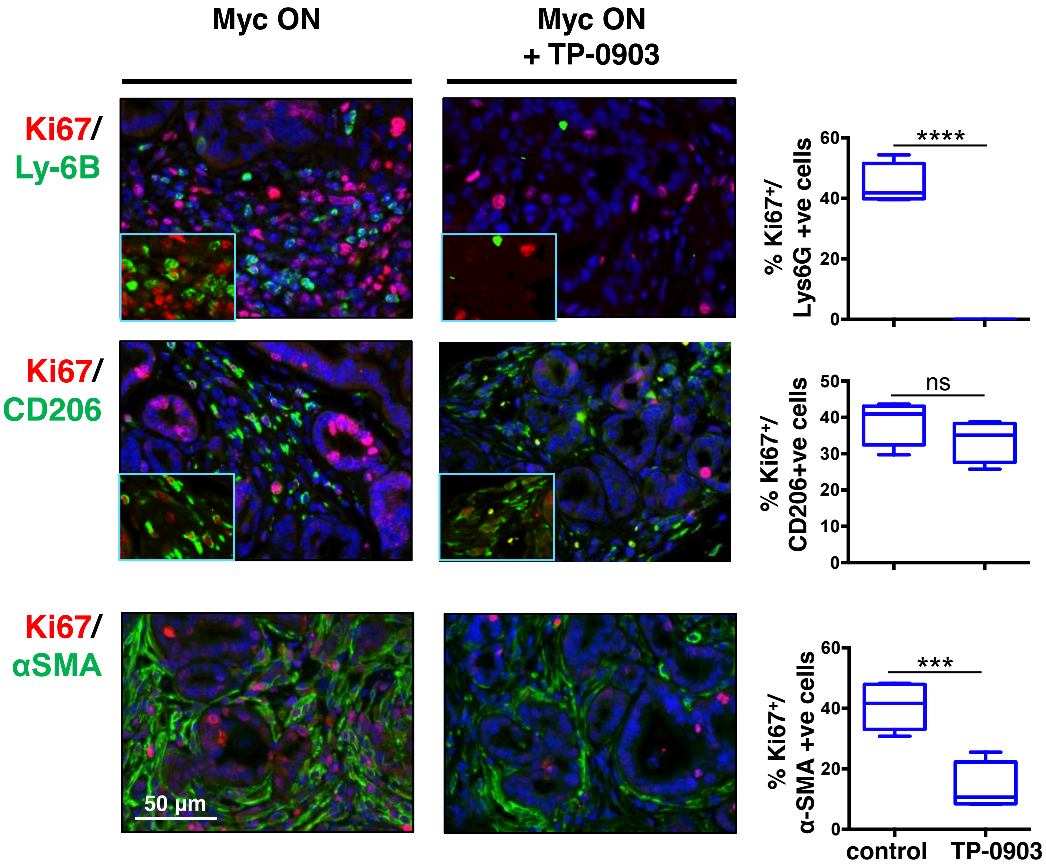

**Figure S6 (related to Figure 5)**

**Myc directly induces *PD-L1* expression in pancreatic epithelial cells**

(A) Immunofluorescence staining and quantification of PD-L1 expression in sections of pancreata harvested from 12-week old $KM^{pdx1}$mice treated with either oil (Myc OFF, 0 day) or tamoxifen (Myc ON) for 1, 3 or 7 days. Inset shows boxed region at higher magnification. n = 4 – 6 per group. *p < 0.05, **p < 0.01, ***p < 0.001. FOV = field of view.

(B) Tumor cells derived from the pancreas of a $KM^{pdx1}$mouse treated with tamoxifen (Myc ON) at 12 weeks of age for 3 weeks were maintained *in vitro* in the presence of 4-hydroxytamoxifen (Myc ON). *PD-L1* mRNA expression was determined in $KM^{pdx1}$ cell line propagated on normal media for 1 week (Myc OFF, 0h) following addition of 4-OHT (Myc ON) for 3, 6, 12 hr and 7 days. The unpaired t-test was used to analyze Taqman expression data. The mean ± S.D. from 3 independent experiments is shown.

(C) ChIP-PCR for Myc binding was performed in the primary cell line described in (b) at 0, 6 and 24 hr after 4-OHT addition (Myc ON). ChIP was performed with rabbit polyclonal anti-cMyc, rabbit polyclonal anti ER$\alpha$ antibodies, or a purified IgG as control. Quantitative PCR for *PD-L1* DNA is expressed as a percentage of the input. The unpaired t-test was used to analyze quantitative expression data. The mean ± SD is shown. n = 3 PCR replicates. *p < 0.05, ***p < 0.001, ns = not significant.

Sodir *et al*. Figure S6 (related to Figure 5)

#### Myc directly induces *PD-L1* expression in pancreatic epithelial cells

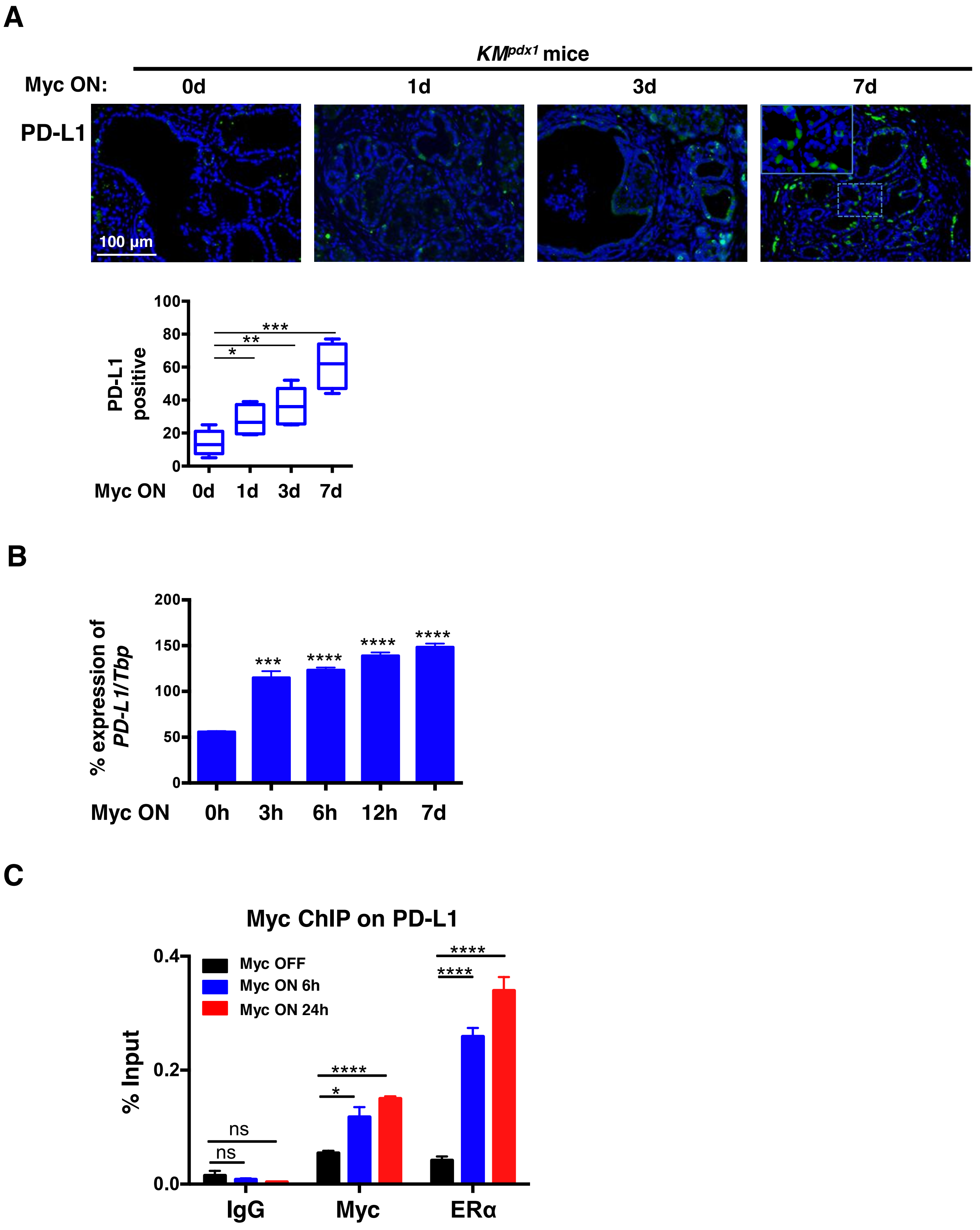

**Figure S7 (related to Figure 6)**

**De-activation of Myc triggers macroscopic regression of *KM*$^{p48}$ pancreatic tumors**

(A) Representative ultrasound imaging of pancreatic tumors in *KM*$^{p48}$ treated with tamoxifen (Myc ON) for 6 weeks at 12 weeks of age (left) and after withdrawal of tamoxifen (Myc OFF) for 3 weeks (middle) or 6 weeks (right). Large cysts (white arrows) form following tamoxifen withdrawal-induced regression. Pancreas (blue line), stomach (red line) and kidney (green line) are all indicated. Tumor sizes were estimated from ultrasound images and are shown on the right.

(B) Representative images of whole pancreas and spleen harvested from *KM*$^{p48}$ mice treated with tamoxifen (Myc ON) for 6 weeks (left panel) or after subsequent Myc deactivation (Myc OFF) for 6 weeks (middle and right panels).

(C) Representative H&E stained sections of *KM*$^{p48}$ pancreata after sustained administration of tamoxifen for 6 weeks (left panel, Myc ON) and then after Myc withdrawal for 6 weeks (middle and right panels.

Sodir *et al*. Figure S7 (related to Figure 6)

#### De-activation of Myc triggers macroscopic regression of *KM*$^{p48}$ pancreatic tumors

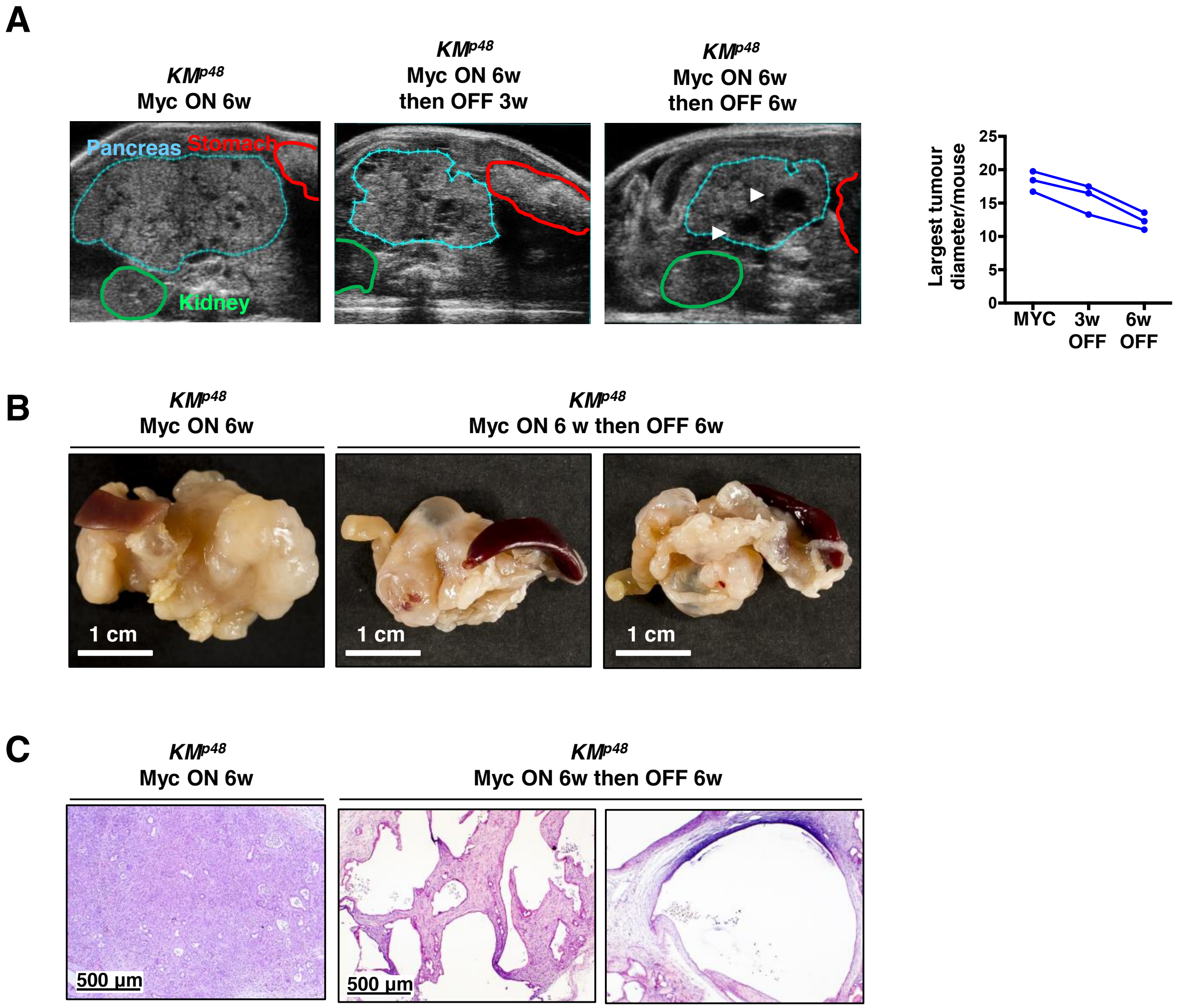

**Figure S8 (related to Figure 7)**

**Myc de-activation triggers immediate reversal of *KM*$^{p48}$ tumor stroma**

Immunohistochemical staining (IHC) for proliferation (Ki67), immunofluorescence (IF) staining of cell death (TUNEL), macrophages (F4/80) and neutrophils (Ly-6B), IHC of T cells (CD3), IF of PD-L1, IHC of B220 lymphocytes and α-SMA, Gomori's Trichrome staining of collagen, and IF for endothelial cells (CD31) and hypoxia (HIF-1α) in sections of pancreata harvested from 12 week old *KM*$^{p48}$ mice treated with tamoxifen (Myc ON) for 3 weeks (0 day, left) followed by tamoxifen withdrawal (Myc OFF) for 3 days (right). Scale bars apply across each row of the panel. Quantification is shown on the right. Data were analyzed using unpaired t- test. n = 4-6 for each treatment group. *p < 0.05, **p < 0.01, ***p < 0.001, ****p < 0.0001 (ns = non-significant). FOV = field of view.

#### Myc de-activation triggers immediate reversal of *KM^p48* tumor stroma

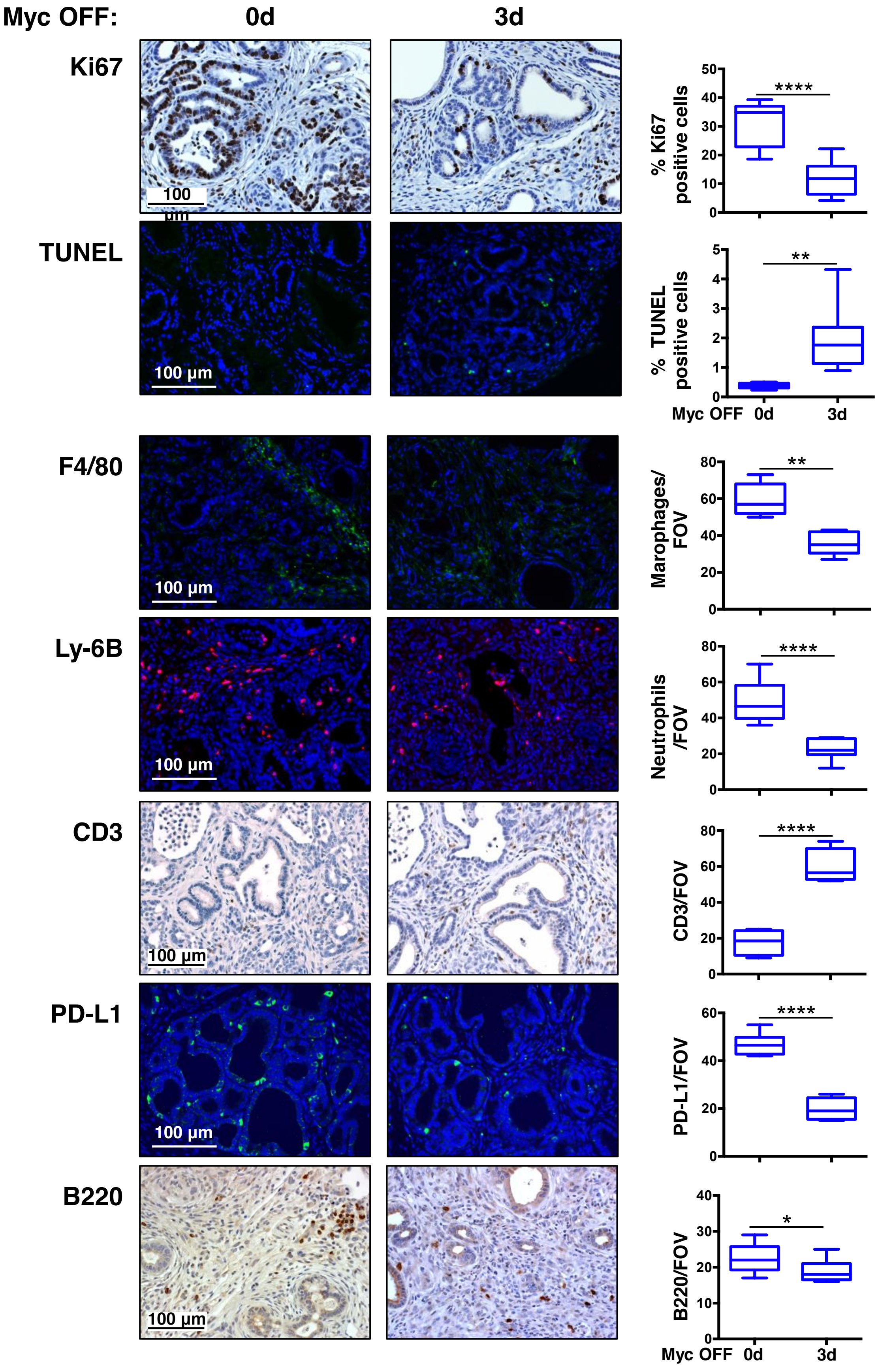

### Myc de-activation triggers immediate reversal of *KM*$^{p48}$ tumor stroma

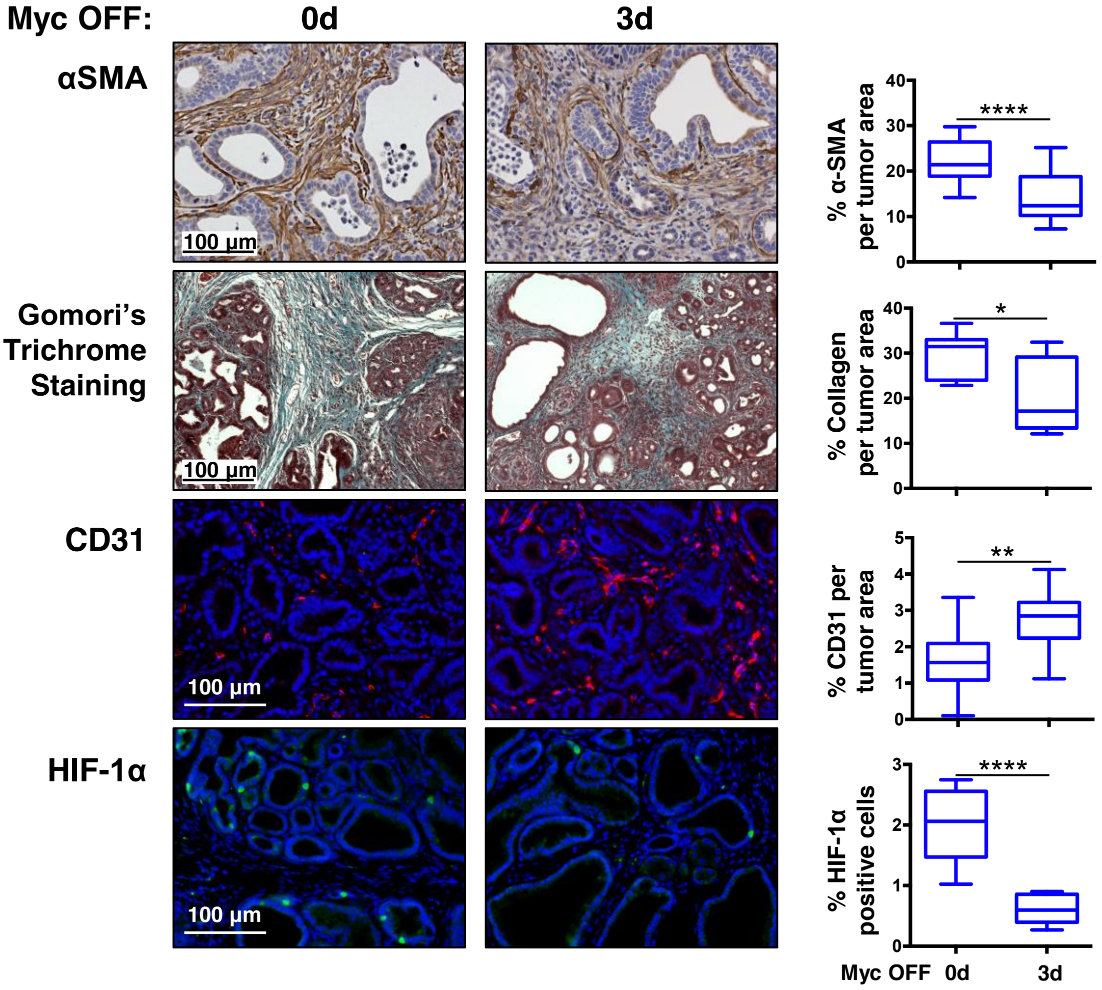

**Table 1**

**Comparison of Myc-induced phenotypes in KRas$^{G12D}$-driven PanIN versus KRas$^{G12D}$-driven lung adenoma**

Sodir *et al.* Table 1

#### Comparison of Myc-induced phenotypes in $KRas^{G12D}$-driven PanIN versus $KRas^{G12D}$-driven lung adenoma

| Pancreas | Lung |
|---|---|
| Increase in epithelial proliferation | |
| Rapid increase in local invasiveness | |
| Immediate macrophage influx | |
| Immediate T cell efflux | |
| Immediate neutrophil influx | No change in neutrophils (already high) |
| Induced trans-differentiation of mesenchymal cells to myofibroblasts | No evident trans-differentiation to myofibroblasts |
| Induction of α-SMA | No Induction of α-SMA |
| Extensive desmoplastic reaction | No desmoplastic reaction |
| Vascular involution | Profound VEGF-driven angiogenesis |
| Normoxia to hypoxia | Hypoxia to normoxia |
| PDL-1 induction on tumor epithelial cells | PDL-1 on incoming macrophages |
| Induction of CCL9, CCL2, CXCL5 and Gas6 expression | Induction of CCL9 and IL12p40/70 expression |
| Increase in pAxl | pAxl not detected |
| Rapid influx in B cells to tumor periphery | Immediate B cell efflux |
| No significant NK cells present | NK cells rapidly expelled from peri-tumoral tertiary lymphoid structures |
